# Visual versus visual-inertial guidance in hawks pursuing terrestrial targets

**DOI:** 10.1101/2022.12.24.521635

**Authors:** James A. Kempton, Caroline H. Brighton, Lydia A. France, Marco KleinHeerenbrink, Sofia Miñano, James Shelton, Graham K. Taylor

**Affiliations:** Department of Biology, University of Oxford, OX1 3SZ, UK

## Abstract

The flight behaviour of predatory birds is well modelled by a guidance law called proportional navigation, which commands steering in proportion to the angular rate of the line-of-sight from predator to prey. The line-of-sight rate is defined with respect to an inertial frame of reference, so proportional navigation is necessarily implemented using visual-inertial sensor fusion. In Harris’ hawks, pursuit of terrestrial targets may be even better modelled by assuming that visual-inertial information on the line-of-sight rate is combined with visual information on the deviation angle between the attacker’s velocity and the line-of-sight. Here we ask whether a new variant of this mixed guidance law can model Harris’ hawk pursuit behaviour successfully using visual information alone. We use high-speed motion capture to record n=228 attack fights from N=4 Harris’ hawks, and confirm that proportional navigation and mixed guidance using visual-inertial information both model the trajectory data well. Moreover, the mixed guidance law still models the data closely if visual-inertial information on the line-of-sight rate is replaced with purely visual information on the apparent motion of the target relative to the background. Although the original form of the mixed guidance law provides the closest fit, all three models provide an adequate phenomenological model of the behavioural data, whilst each making different predictions on the physiological pathways involved.

## Introduction

A hawk pursuing a hare must steer its attack trajectory to achieve capture. Drawing inspiration from missile engineering, an increasingly popular approach to understanding such behaviours is to identify a behavioural algorithm, or guidance law, that successfully models the turning behaviour of the attacker (***Hein et al., 2020***). Guidance laws implement pursuit by commanding turning in response to the relative motion of an attacker and its target (***Shneydor, 2011***). For example, a guidance law called proportional navigation (PN) has long been used to implement guidance in homing missiles (***Palumbo et al., 2010***), and commands steering in proportion to the angular rate 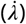 of the line-of-sight from the attacker to its target. An alternative guidance law called proportional pursuit (PP) involves commanding turning in proportion to the deviation angle (*δ*) between the attacker’s velocity and the line-of-sight from the attacker to its target. These different guidance laws predict different pursuit trajectories in response to the same target motion, and whereas PN guidance closely models the steering behaviour of falcons (***Brighton et al., 2017***, ***2021***), bats (***Ghose et al., 2006***), robber flies (***Fabian et al., 2018***), and killer flies (***Fabian et al., 2018***), PP guidance has been found to model the steering of blowflies (***Boeddeker and Egelhaaf, 2003***), tiger beetles (***Haselsteiner et al., 2014***) and honeybees (***Zhang et al., 1990***). Other, more complex, guidance laws are possible. For example, ***Brighton and Taylor*** (***2019***) found that combining PN and PP guidance linearly in a mixed (PNP) guidance law most successfully modelled the steering behaviour of Harris’ hawks (*Parabuteo unicinctus*), which are the subject of this study.

Besides commanding different pursuit trajectories in response to the same target motion, the PN, PP, and PNP guidance laws differ fundamentally in the nature of the sensory cues they use. PN commands steering in response to the line-of-sight rate, 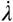, defined as the rate of change in the line-of-sight angle (*λ*) measured between the line-of-sight to the target and some fixed reference direction (Figure 1). It follows that 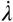 is an exocentric cue, because measuring it involves measuring changes in the line-of-sight direction with respect to an external frame of reference. In contrast, PP commands steering in response to the deviation angle, *δ*, defined as the angle between the line-of-sight to the target and the attacker’s own velocity vector (Figure 1). In principle, *δ* can be thought of as an egocentric cue of sorts, because it involves measuring the line-of-sight direction with respect to the attacker’s own self-motion. In homing missiles, at least, this can make PP guidance simpler to mechanise than PN (***Shneydor, 2011***). Finally, the mixed PNP guidance law combines PN and PP elements, and therefore employs a mixture of ego- and exocentric cues. Nevertheless, as we show below, a related form of the same mixed guidance law can be made to work using egocentric cues alone. As we now discuss, this distinction is important because of the difference it makes to how the respective variables can be measured physiologically.

**Figure 1.**
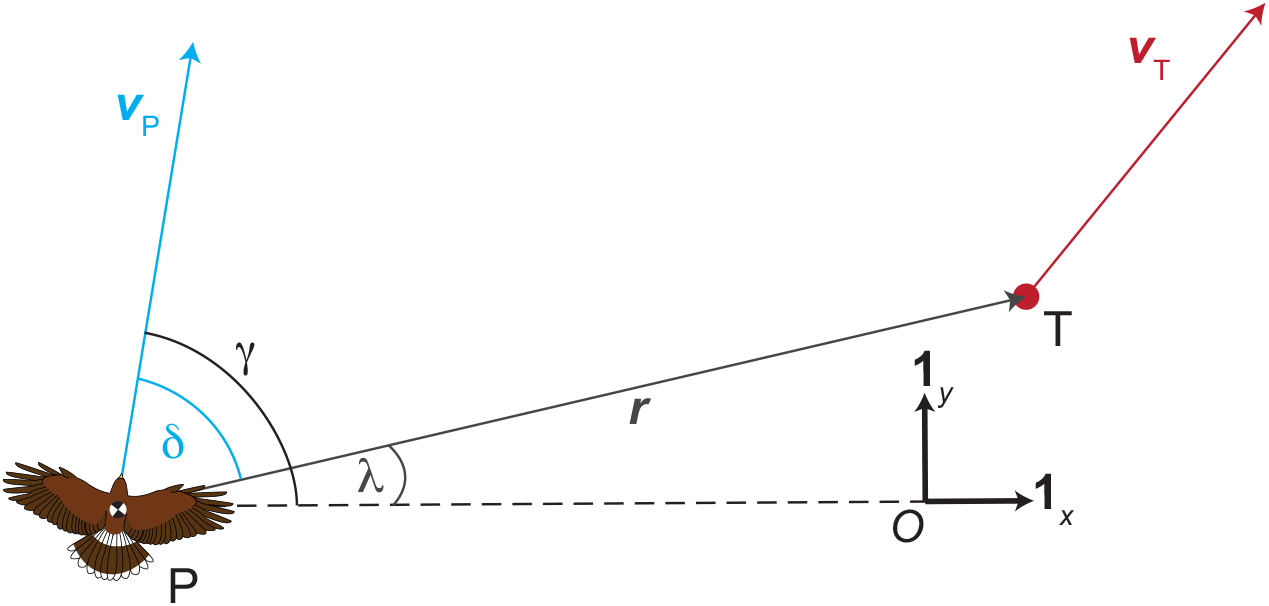
Pursuit geometry. The fixed orthonormal basis vectors **1_*x*_** and **1_*y*_** are taken to define an inertial frame of reference 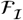 in which the various quantities are measured. The angle between the hawk’s velocity vector ***v_P_*** and the inertial reference direction **1_*x*_** is the flight path angle *γ*. The line drawn from the pursuer (P) to its target (T) is called the line-of-sight vector ***r***. The angle between the line-of-sight vector ***r*** and the inertial reference direction **1_*x*_** is called the line-of-sight angle *λ*. The time derivative of the line-of-sight angle with respect to 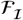 is called the inertial line-of-sight rate 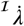, and is fed back to command turning under the inertial-PN guidance law. The angle that the hawk’s velocity vector, ***v_P_***, forms with the line-of-sight vector ***r*** is the target deviation angle, *δ*, which is fed back to command turning under the proportional pursuit (PP) guidance law. The inertial-PNP guidance law, which has been used to model Harris’ hawk attack trajectories previously (***Brighton and Taylor, 2019***), feeds back both 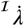 and *δ* to command turning.

In homing missiles (***Palumbo et al., 2010***; ***Shneydor, 2011***), and in studies of animal pursuit (***Brighton et al., 2017***, ***2021***; ***Fabian et al., 2018***), the line-of-sight angle *λ* is conventionally defined with respect to a fixed reference direction. In practice, this means that any agent which employs, or is hypothesised to employ, PN guidance must be able to measure the line-of-sight rate 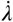 in an inertial frame of reference, or in some other fixed frame of reference approximating this. A typical homing missile does so by rotating its target acquisition sensor (e.g. radar dish or imaging sensor) to centre the target, using a gimbaled tracking device called a seeker. Rate gyros sensing the seeker’s angular velocity in an inertial frame of reference are then used to estimate 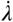 as the rotation rate of the seeker minus any residual tracking error. Although birds may use head movements to track targets visually in a similar fashion to a missile seeker, it has not yet been tested whether or how they estimate the line-of-sight rate 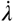. It is likely the sensory output of the vestibular system could be used for this purpose, but whilst this is well set up to sense angular acceleration of the head, it remains uncertain whether the vestibular system can measure the angular rate of the head directly without integration and hence drift (***Warrick, 2002***; ***Pennycuick, 2008***; ***Muller, 1990***, ***2020***).

Although it is reasonable to suppose that vestibular output may be used to estimate the tracking rate of the head in response to target motion, the hypothesis that the head functions similarly to a missile seeker already assumes that it tracks the target continuously in flight. In practice, a bird’s head is also used to look around the environment when acquiring a target, and is typically turned in a discontinuous fashion, with longer periods of stabilized gaze interspersed with brief saccadic motions. It follows that vestibular output may instead be used to stabilize the head in an inertial frame of reference (***Benson et al., 2017***), whence the visual drift of the target across the retina would provide a direct measure of the inertial line-of-sight rate 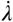. Finally, vision can provide an inertial reference if there are landmarks present that do not change appreciably with the attacker’s self-motion. It is even possible that the sun’s disc or its attendant cues could provide such a reference (***Guilford and Taylor, 2014***). However, a hawk chasing a target close to the ground or beneath the canopy will see no distant background against which to measure the target’s motion, and the direction of nearby visual features will change quickly as a result of its own self-motion. It therefore seems reasonable to assume that animals implementing classical PN guidance will do so with the aid of inertial sensors, used either to measure the rotation rate of their head or to stabilize their head rotationally.

In contrast to the line-of-sight rate 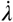, the deviation angle *δ* can be measured without an inertial reference. For example, if the background is close enough to provide parallax cues, then the deviation angle *δ* can be measured visually, by taking the vanishing point of the translational optic flow field as the direction of self-motion, and comparing this to the retinal coordinates of the target. Alternatively, the deviation angle could be estimated from the contralateral asymmetry in the translational optic flow field (***Ros and Biewener, 2017***) that is produced when a predatory bird looks directly at its target. Hence, the hypothesis that an organism implements PP guidance makes fundamentally different assumptions regarding its sensory capabilities as compared to the hypothesis that an organism implements PN guidance. Clearly, the combination of PN and PP elements in PNP guidance rests on a different set of assumptions again. It follows that phenomenological modelling of pursuit behaviour can in principle be informative as to the physiological pathways that may be involved. For instance, ***Brighton and Taylor (2019***) found that whereas PNP or PN guidance could both successfully model Harris’ hawk attack trajectories, PP guidance could not. Given the necessary involvement of a PN element, this result would seem to imply that hawks must be able to measure the line-of-sight rate 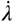 in an inertial reference frame. However, as we now show, other mixed guidance models that do not imply the ability to measure 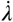 in an inertial frame of reference may model Harris’ hawk trajectories as well as, or even better than, the PNP guidance law that ***Brighton and Taylor (2019***) fitted.

Here we describe a new data set comprising n=228 unobstructed pursuit trajectories flown by N=4 Harris’ hawks. The same data are used as baseline data elsewhere in an analysis of obstacle avoidance in obstructed pursuit ***Brighton et al. (2023***) utilising a slightly different fitting method (see Methods and Materials). Here we model these unobstructed pursuit data using the original PP, PN, and PNP guidance laws described above, together with new versions of the PN and PNP guidance laws in which the line-of-sight rate is defined with respect to the visual background behind the target. This background line-of-sight rate, which we write as 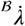, is defined as the difference between the retinal drift of the target and the retinal drift of the background. It can therefore be measured without the involvement of inertial sensors or an inertial reference, in contrast to the inertial line-of-sight rate, which we write as 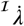 for clarity. Having reframed PN and PNP guidance by substituting 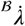 in place of 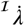, we test the success of the resulting background-PN and background-PNP guidance laws in explaining the measured hawk trajectories relative to inertial-PN and inertial-PNP. Although background-PN performs poorly for reasons we explain below, we find that background-PNP explains our data at least as effectively as inertial-PN, and almost as well as inertial-PNP. This has important implications for the sensorimotor pathways that are plausibly involved in implementing pursuit guidance in hawks and other flying animals, since background-PNP can be implemented without necessarily involving inertial sensors in the guidance loop.

## Theoretical framework

Figure 1 shows the geometry of a pursuit involving a hawk chasing a terrestrial target over a flat horizontal surface. We will neglect any vertical displacement of the pursuer, which is assumed to be small in comparison to its horizontal displacement. The line-of-sight vector ***r*** is defined as the position vector of the target with respect to the pursuer in an external frame of reference defined by the orthonormal basis vectors **1_*x*_** and **1_*y*_**. The pursuer’s flight path angle *γ* is defined as the signed angle that its velocity vector ***v_P_*** makes with the external reference direction **1_x_**. Meanwhile, the line-of-sight angle *λ* is defined as the signed angle that the line-of-sight vector ***r*** makes with the reference direction **1_x_**, and the deviation angle *δ* is defined as the signed angle that the line-of-sight vector ***r*** makes with the pursuer’s velocity vector ***v_P_***.

It is clear from Figure 1 that the deviation angle *δ* is independent of the choice of external reference direction **1_*x*_**, so can in principle be measured egocentrically. In contrast, both the flight path angle *γ* and the line-of-sight angle *λ* depend on the choice of external reference direction **1_*x*_**, and must therefore be measured exocentrically. To begin with, we will fix **1*_x_*** and **1_*y*_** so that they define an inertial frame of reference 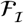, and will use the notation 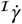 and 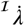 to denote the time derivatives of *γ* and *λ* in 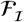. With these definitions, the proportional pursuit (PP) guidance law commands turning as:

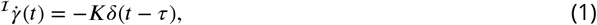

where *K* is a guidance constant and *τ* is a fixed time delay. In contrast, the proportional navigation (PN) guidance law commands turning as:

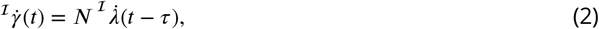

where *N* is another guidance constant. Finally, the PNP guidance law that ***Brighton and Taylor*** (***2019***) defined commands turning as:

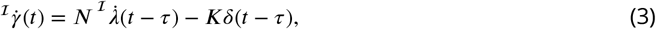

where, for sake of simplicity, the same time delay *τ* is assumed to hold for both elements.

Because we have fixed **1_*x*_** and **1_*y*_** with respect to an inertial frame of reference, practical methods of estimating the inertial line-of-sight rate 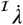 are expected to involve the fusion of visual and inertial information. One possibility is to track the target visually by turning the head, and to measure the angular rate of the head inertially as a proxy for 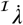. An alternative is to stabilise the head inertially, and to measure the retinal drift of the target visually as a proxy for 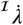. However, we may choose to use other sets of basis vectors to define the line-of-sight rate, and in this case the sensory basis of its measurement may differ. For instance, we may choose to define a new external reference direction **1_*x*′_** visually (e.g. as the direction vector of some background feature seen by the pursuer), and may use **1_*x*′_** and its normal **1_*y*′_** to define a new background frame of reference 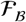. If the background feature used to define **1_*x*′_** is sufficiently distant (e.g. the sun’s disc), then 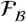 will approximate an inertial frame of reference. This being so, it is possible to estimate 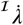 visually by comparing the retinal drift of the target to the retinal drift of the distant background. On the other hand, if the background feature is close to the pursuer (e.g. a patch of ground beneath the target), then 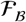 will rotate as the pursuer moves. In this case, the difference between the retinal drift of the target and the retinal drift of the background is no longer a proxy for the inertial line-of-sight rate 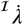, but still measures a well-defined quantity that we will call the background line-of-sight rate 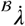.

In summary, we therefore have one version of the line-of-sight rate 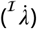 that is defined in an inertial frame of reference 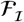, and another 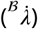 that is defined with respect to the background frame of reference 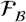. In computing the background line-of-sight rate 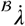, we will assume that the distance between the target and the background feature used to define 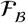 is negligibly small, which is reasonable in the case of a hawk chasing a terrestrial target moving over the ground. With this restriction, the background line-of-sight rate 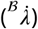 describes only the component of the relative motion of the target and pursuer that is due to the target’s motion (see Supplementary Text). In contrast, the inertial line-of-sight rate 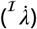 describes the total motion of the target relative to the pursuer, so also contains information on the pursuer’s own self-motion. Substituting 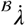 for 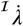 in the PN guidance law therefore loses information on self-motion that is useful for steering an effective intercept. Thus, the following guidance law, which we call background-PN to distinguish it from the inertial form of the PN guidance law described above, is unlikely to steer attack trajectories successfully:

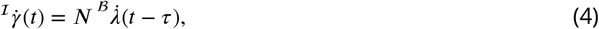

However, as we show in the Supplementary Text, substituting 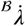 for 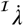 in the mixed PNP guidance law results in no net loss of information, because of the complementary effect of the deviation angle *δ*. With this in mind, we define the following new guidance law, which commands steering as:

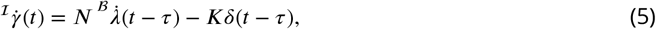

which we call the background-PNP guidance law to distinguish it from the inertial form of the PNP guidance law described above. Here we ask whether background-PNP can model Harris’ hawk attack trajectories as successfully as inertial-PNP, and compare this to the performance of the inertial-PN, background-PN, and PP guidance laws in modelling the same data.

## Results

### Guidance model simulations

We recorded N=4 Harris’ hawks pursuing an artificial lure being pulled around a set of pulleys on n=228 trials, and used a motion capture system to measure the trajectories of the hawk and target at 200 Hz. For each trial, we simulated the horizontal components of the hawk’s measured flight trajectory under the assumption that its steering was modelled by any one of the five candidate guidance laws in Table 1. We matched the hawk’s simulated flight speed to its measured flight speed, which allowed us to simulate its trajectory given knowledge of the hawk’s initial position and measured ground speed, together with knowledge of the lure’s trajectory throughout the trial (see Methods and Materials for details). This is the same modelling approach taken in our previous work (***Brighton et al., 2017***; ***Brighton and Taylor, 2019***; ***Brighton et al., 2021***), but with the addition of two new candidate guidance laws in the form of background-PN and background-PNP. We begin by fitting the parameters of these five guidance models independently to each flight, before then fitting the same parameters globally to all flights to guard against over-fitting. Throughout the paper, we use hat notation (e.g. 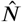) to denote a parameter estimate for a given flight, tilde notation (e.g. 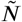) to denote the median of these parameter estimates over all n=228 flights, and breve notation (e.g. 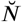) to denote the globally-fitted values of the model parameters. We report two-tailed test statistics throughout, and treat each flight as an independent sample noting that the sample sizes we state represent technical (n=228) rather than biological (N=4) replicates.

**Table 1.** Guidance laws fitted to the data, together with the equations describing them, and the sensory information they imply that a hawk would require to implement a pursuit.

| Model | Equation | Information required |
| --- | --- | --- |
| PP | ${}^I\dot{\gamma}(t) = -K\delta(t - \tau)$ | visual-only |
| inertial-PN | ${}^I\dot{\gamma}(t) = N^I\dot{\lambda}(t - \tau)$ | visual-inertial |
| background-PN | ${}^I\dot{\gamma}(t) = N^B\dot{\lambda}(t - \tau)$ | visual-only |
| inertial-PNP | ${}^I\dot{\gamma}(t) = N^I\dot{\lambda}(t - \tau) - K\delta(t - \tau)$ | visual-inertial |
| background-PNP | ${}^I\dot{\gamma}(t) = N^B\dot{\lambda}(t - \tau) - K\delta(t - \tau)$ | visual-only |

### Guidance models fitted independently to each flight

We begin by fitting the parameters *N*, *K*, and *τ* for each flight independently under the five candidate guidance models in Table 1. We do so by finding the parameter values 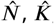, and 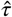 that minimise the root mean square (RMS) error (*ϵ*) between the measured and simulated trajectories for each flight (Figure 3). The results of the model fitting are summarised in Table 2 and Figure 4, which show that the two alternative forms of the PNP guidance law are both capable of fitting the data closely. In fact, the median RMS errors under inertial-PNP (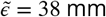; 95% bootstrapped CI: 28, 51 mm) and background-PNP (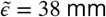; 95% bootstrapped CI: 29, 51 mm) were statistically indistinguishable over the n=228 flights (sign test: *p* = 0.64). Nevertheless, the inertial-PNP and background-PNP models are not equivalent, as can be seen from the fact that the best-fitting models under these two alternative forms of PNP guidance generate different simulated trajectories for the example flight plotted in Figure 3. But how, exactly, does this difference arise?

**Figure 2.**
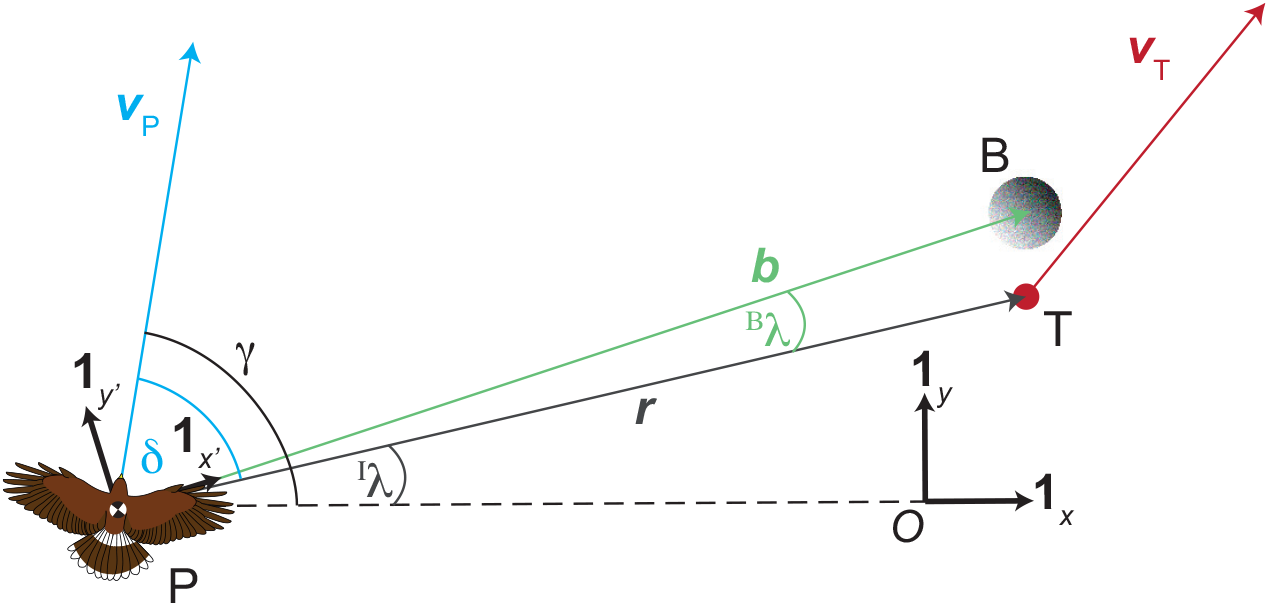
Harris’ hawk attack geometry with background. Here we add to the geometry of Figure 1 a background feature (B) that lies close to the target (T). We use the rotating orthonormal basis vectors 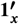 and 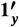 carried with the bird to define the background reference frame 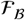, where 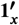 points in the direction of the vector ***b*** from the hawk (P) to the background feature (B). Note that 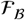 is defined instantaneously by the position of the hawk relative to the background patch against which it is viewing the target. As such, the background reference frame 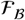 is constantly updated, in contrast to the inertial reference frame 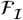 which is fixed for all time. Rewriting the line-of-sight angle *λ* as *^I^λ* shows that this is measured with respect to the inertial reference direction **1_x_**, in contrast to the background line-of-sight angle *^B^λ* which is measured with respect to the background reference direction 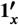. The time derivative of the line-of-sight angle with respect to 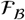 is called the background line-of-sight rate *^B^λ*, and is fed back to command turning under the background-PN and background-PNP guidance laws.

**Figure 3.**
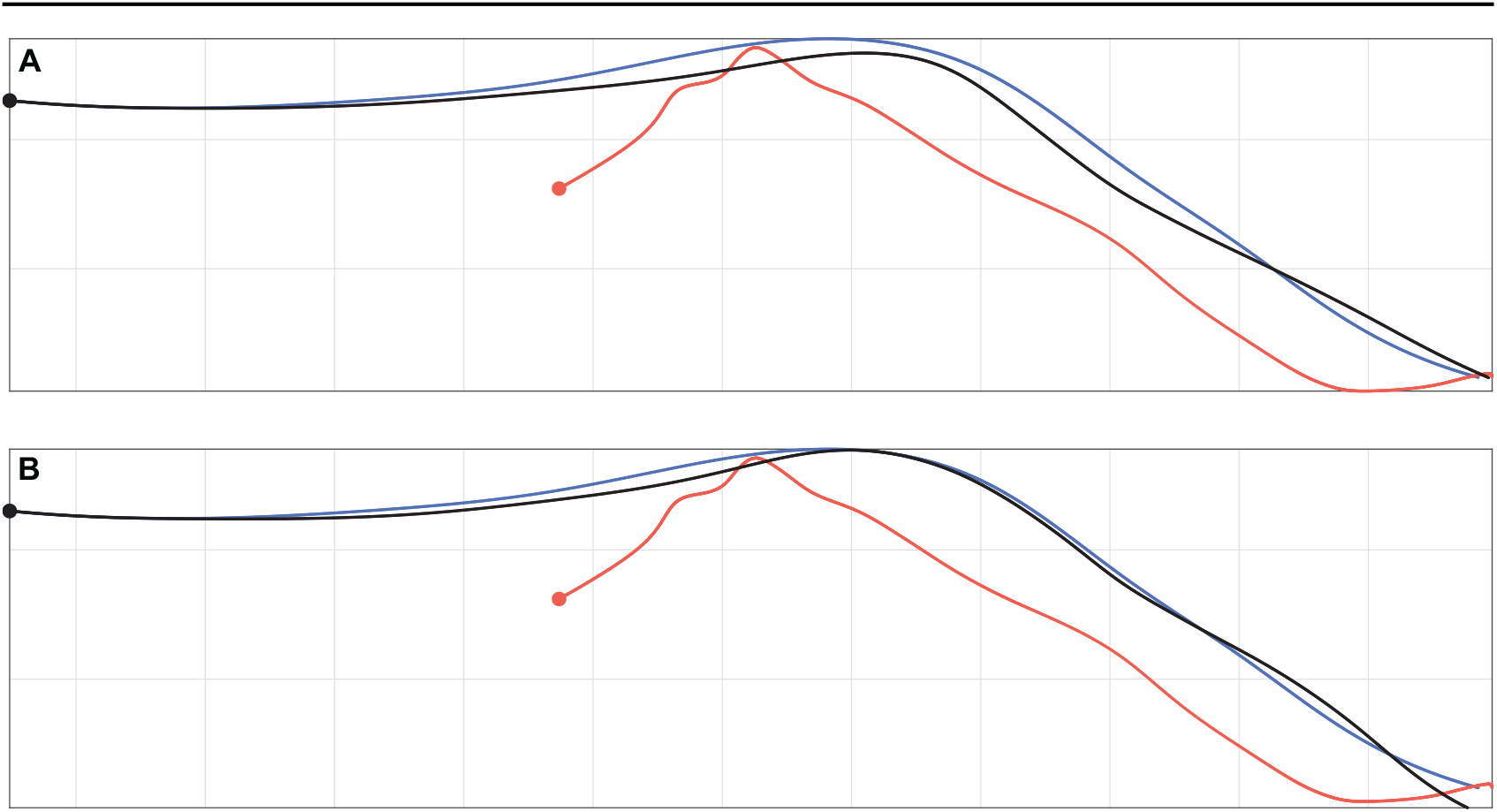
Measured trajectory for one example flight compared to simulations under inertial-PNP and background-PNP. Given the measured trajectories of the hawk (blue) and lure (red), we generate the hawk’s simulated trajectory (black) by solving for the guidance model parameters that minimise the RMS error 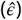 between the measured and simulated trajectories. **A** Best-fitting simulation under inertial-PNP, which assumes that the line-of-sight rate is measured in an inertial frame of reference 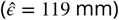; **B** Best-fitting simulation under background-PNP, which assumes that the line-of-sight rate is measured in the background frame of reference 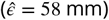. Note that in this particular example, the background-PNP model fits the measured data more closely than the inertial-PNP model; in other examples, this situation may be reversed. Grid size: 1 m.

**Table 2.** Summary of the results of fitting the five alternative guidance models to each flight independently.

| Guidance model | Median (95% CI) of RMS error (mm) | Median (95% CI) of best-fitting guidance parameters |
| --- | --- | --- |
| PP | $\tilde{\epsilon} = 135$ (111, 154) | $\tilde{K} = 4.4 \text{ s}^{-1}$ (3.2, 6.1)<br>$\tilde{\tau} = 0.030 \text{ s}$ (0.015, 0.045) |
| inertial-PN | $\tilde{\epsilon} = 71$ (54, 90) | $\tilde{N} = 0.85$ (0.77, 0.91)<br>$\tilde{\tau} = 0.060 \text{ s}$ (0.050, 0.068) |
| background-PN | $\tilde{\epsilon} = 145$ (113, 203) | $\tilde{N} = 0.66$ (0.51, 0.82)<br>$\tilde{\tau} = 0.073 \text{ s}$ (0.050, 0.090) |
| inertial-PNP | $\tilde{\epsilon} = 38$ (28, 51) | $\tilde{N} = 0.77$ (0.66, 0.86)<br>$\tilde{K} = 0.9 \text{ s}^{-1}$ (0.51, 1.4)<br>$\tilde{\tau} = 0.058 \text{ s}$ (0.040, 0.065) |
| background-PNP | $\tilde{\epsilon} = 38$ (29, 51) | $\tilde{N} = 0.66$ (0.51, 0.82)<br>$\tilde{K} = 3.4 \text{ s}^{-1}$ (2.8, 3.8)<br>$\tilde{\tau} = 0.060 \text{ s}$ (0.048, 0.070) |

**Figure 4.**
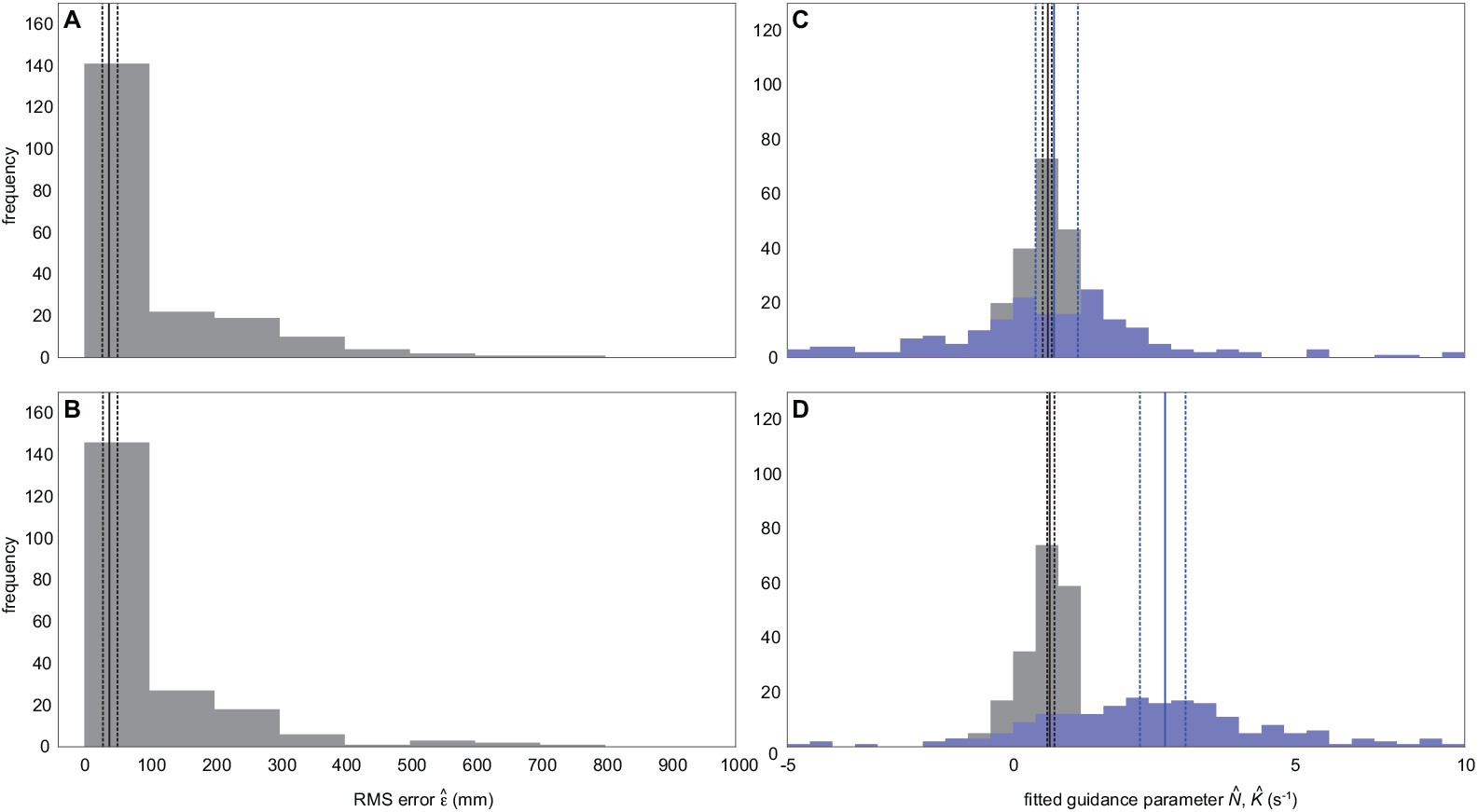
Distributions of the RMS error and guidance parameters fitted to each flight under inertial-PNP and background-PNP. Histograms of **(A,B)** the RMS error 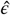 (grey), and **(C,D)** the guidance parameters 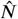 (grey) and 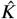 (blue), for the best-fitting guidance models under inertial-PNP (A,C) and background-PNP (B,D) when fitted to each flight independently. A solid vertical line denotes the median of each distribution; broken vertical lines display the corresponding bootstrapped 95% confidence interval.

The median best-fitting values of *N* were similar under inertial-PNP (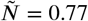; 95% bootstrapped CI: 0.66, 0.86) and background-PNP (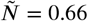; 95% bootstrapped CI: 0.51, 0.82), and their respective confidence intervals each contain the best-fitting value of *N* = 0.7 that ***Brighton and Taylor*** (***2019***) identified under inertial-PNP in their previous study of Harris’ hawks. Likewise, the median best-fitting value of *K* under inertial-PNP (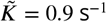; 95% bootstrapped CI: 0.51, 1.4 s^−1^) was statistically indistinguishable from the best-fitting value of *K* = 1.0 s^−1^ that ***Brighton and Taylor*** (***2019***) identified under inertial-PNP. On the other hand, the best-fitting values of *K* under background-PNP were significantly higher than those under inertial-PNP (sign test: *p* < 0.0001; n=228 flights), and the confidence interval for their median (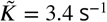; 95% bootstrapped CI: 2.8, 3.8 s−1) does not contain the best-fitting value of *K* = 1.0 s^−1^ identified under inertial-PNP by ***Brighton and Taylor*** (***2019***). This is a striking result, because whereas the parameter *K* describes how the pursuer responds to the target deviation angle, which is defined in the same way in both models, the parameter *N* describes how the pursuer responds to the target line-of-sight rate, which is defined differently under inertial-PNP and background-PNP. Yet, it is the fitted values of the parameter *K* which are 199 sensitive to the definition of the line-of-sight rate; a result which we explain fully in the Discussion.

In any case, the spread of the fitted values of *K* under both forms of PNP is suggestive of over-fitting (Figure 4). This evidence of over-fitting is particularly clear for inertial-PNP, since the interquartile range of the fitted values of *K* spans zero (Figure 4), which means that even the sign of this guidance parameter estimate is inconsistent between flights. Moreover, although inertial-PN fits the data significantly less well than inertial-PNP over the n=228 flights (sign test: *p* < 0.0001), which reflects the fact that inertial-PN is just a special case of inertial-PNP with *K* = 0, it is clear by inspection of Table 2 that inertial-PN is still capable of fitting the data closely (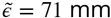; 95% bootstrapped CI: 54, 90 mm). On the other hand, PP and background-PN, which have the same number of fitted parameters as inertial-PN, fit the data significantly less closely than inertial-PN (sign test: *p* < 0.0001; n=228 flights), allowing us to reject them conclusively as models of the data.

In conclusion, we are left with the unequivocal result that inertial-PN (requiring visual-inertial information) models the data much better than either PP or background-PN (for which only visual information is required). We also have the somewhat equivocal result that inertial-PNP (requiring visual-inertial information) and background-PNP (requiring visual-only information) can each fit the data even more closely than inertial-PN, albeit at the risk of over-fitting, and with no way of distinguishing from this analysis which of inertial-PNP and background-PNP is the better model. It follows that whereas visual-inertial information is necessary to explain the data if only the line-of-sight rate is used to command steering (i.e. under PN), visual-only information may be sufficient if information on the deviation angle is added to information on the line-of-sight rate (i.e. under PNP).

### Guidance models fitted globally to all flights

The results of the previous section show that inertial-PN with two fitted guidance parameters per flight (i.e. 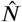 and 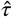), provides a reasonably close fit to the data, but a less close fit than either inertial-PNP or background-PNP with three fitted parameters per flight (i.e. 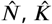, and 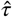). Since the number of fitted parameters scales linearly with the number of flights in this case, it is challenging to interpret this difference from a model selection perspective owing to the risk of over-fitting. In this section, we therefore follow ***Brighton and Taylor*** (***2019***) in taking the complementary approach of fitting a single set of parameters under each guidance model to all n=228 flights, such that we fit only two parameters in total for PP, inertial-PN, and background-PN, and only three parameters in total for inertial-PNP and background-PNP. Specifically, we find the unique parameter values of *γ* and *N* and/or *K* that minimise the median RMS error 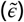 under each of the five guidance models. In practice, because we perform this optimization using an exhaustive search procedure, the global optima for PP, inertial-PN, and background-PN are contained within the optimizations of inertial-PNP and background-PNP, noting that PN and PP are just special cases of PNP with *K* = 0 and *N* = 0, respectively. We therefore plot 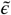 as a function of *N*, *K*, and *τ* under inertial-PNP and background PNP in Figure 5, and summarise the global optimum for each guidance law in Table 3.

**Figure 5.**
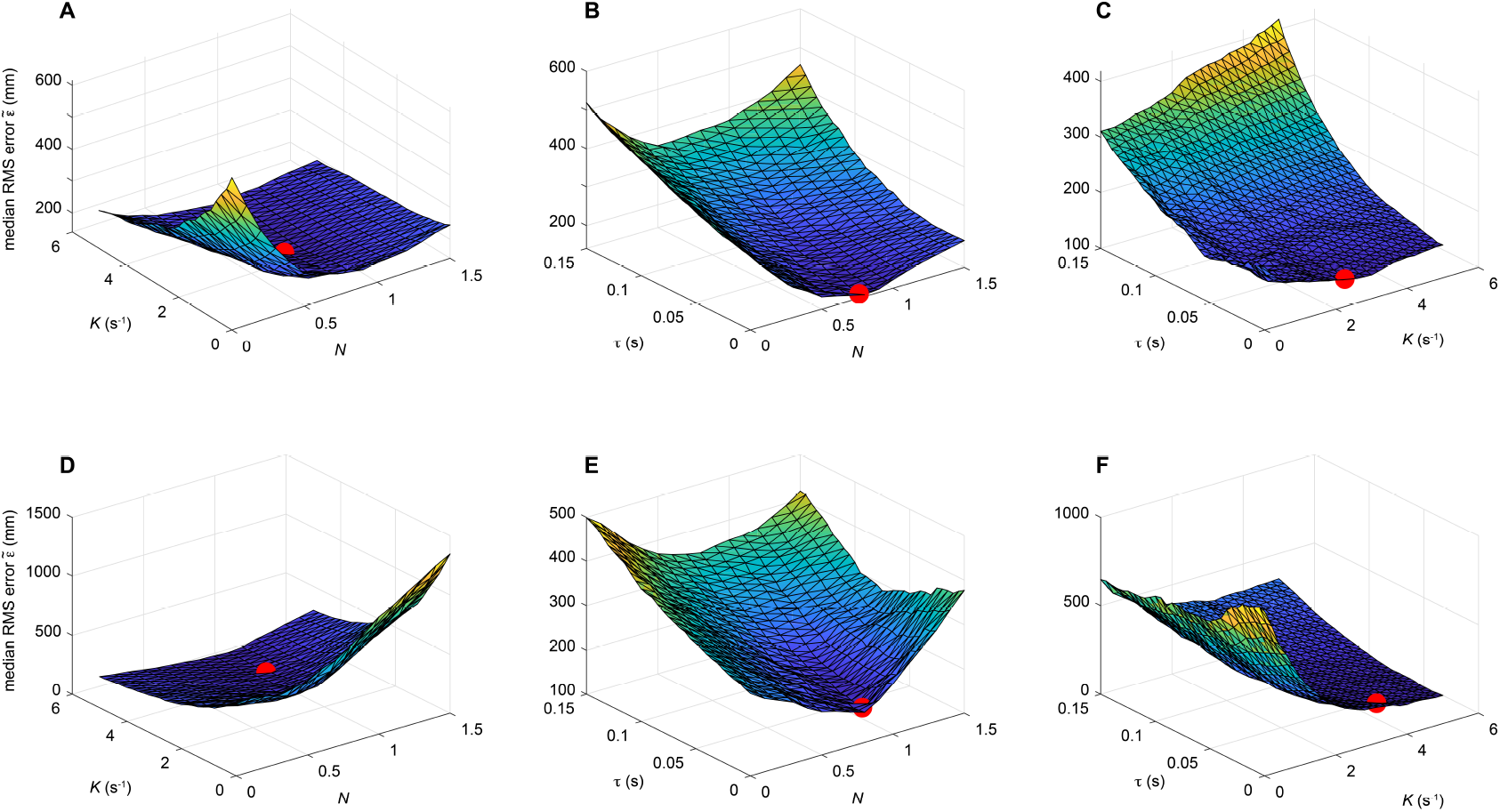
Optimisation of the inertial-PNP and background-PNP models fitted globally to all flights. The surface plots display the median RMS error, 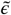, under inertial-PNP **(A-C)** and background-PNP **(D-F)** as a function of the three guidance parameters *N*, *K*, and *τ*, where we vary two parameters at a time and hold the third parameter constant at its best-fitting value. Surface colouring represents 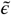; the red spot indicates the optimal parameter combination that minimises 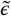 over all flights.

**Table 3.** Summary of the results of fitting the five alternative guidance models globally to all flights.

| Model | Median (95% CI) of RMS error (mm) | Globally optimized values (95% CI) of guidance parameters (SI units) |
| --- | --- | --- |
| PP | $\bar{\epsilon} = 255$ (224, 277) | $\check{K} = 5.0$ (4.8, 5.0)<br>$\check{\tau} = 0$ s (0, 0) |
| inertial-PN | $\bar{\epsilon} = 174$ (148, 225) | $\check{N} = 1.0$ (0.9, 1.1)<br>$\check{\tau} = 0.055$ s (0.035, 0.075) |
| background-PN | $\bar{\epsilon} = 523$ (459, 636) | $\check{N} = 0.5$ (0.3, 0.8)<br>$\check{\tau} = 0.105$ s (0.065, 0.150) |
| inertial-PNP | $\bar{\epsilon} = 141$ (123, 189) | $\check{N} = 0.8$ (0.5, 0.9)<br>$\check{K} = 2.4$ (0.6, 3.6)<br>$\check{\tau} = 0.005$ s (0, 0.030) |
| background-PNP | $\bar{\epsilon} = 150$ (126, 187) | $\check{N} = 0.9$ (0.7, 1.0)<br>$\check{K} = 3.6$ (3.0, 3.8)<br>$\check{\tau} = 0.015$ s (0, 0.045) |

Fitting the guidance models globally to all flights (Table 3) corroborates and extends the results obtained by fitting the guidance models independently to each flight (Table 2). Specifically, the goodness of fit of the guidance models is still similar under inertial-PNP (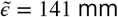; 95% bootstrapped 238 CI: 123, 189 mm) and background-PNP (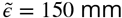; 95% bootstrapped CI: 126, 187 mm), and the difference in their RMS errors remains statistically insignificant over the n=228 flights (sign test: *p* = 0.55). Moreover, neither inertial-PNP (sign test: *p* = 0.44) nor background-PNP (sign test: *p* = 1.0) provides a significantly closer fit than inertial-PN (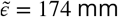; 95% bootstrapped CI: 148, 225 mm) when their model parameters are fitted globally to all flights (Table 3). In contrast, globally-fitted PP displays comparatively poor goodness of fit (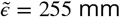; 95% bootstrapped CI: 224, 277 mm) modelling the data significantly less well than inertial-PN (sign test: *p* = 0.007). Globally-fitted background-PN is even less successful at modelling the data, with a very high median RMS error indeed (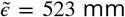; 95% bootstrapped CI: 459, 636 mm). We conclude that the steering of the n=228 Harris’ hawk attack trajectories is comparably well-modelled by inertial-PNP, background-PNP and inertial-PN when the model parameters are fitted globally to all flights. It follows that inertial-PN provides the most parsimonious model assuming use of visual-inertial information, but that background-PNP provides a reasonable model assuming use of visual information only.

The parameter estimates for the PNP guidance models fitted globally to all flights further corroborate and extend the results obtained by fitting those same guidance models independently to each flight. Specifically, the parameter estimates for *N* are similar under globally-fitted inertial-PNP (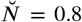; 95% bootstrapped CI: 0.5, 0.9) and globally-fitted background-PNP (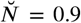; 95% bootstrapped CI: 0.7, 1.0), which in turn are close to the median values of the parameter estimates for *N* under independently-fitted inertial-PNP 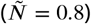 and independently-fitted background-PNP 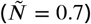. Likewise, the parameter estimate for *K* under globally-fitted background-PNP (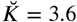 s^−1^; 95% bootstrapped CI: 3.0, 3.8 s^−1^) is close to the median value of the parameter estimate for *K* under independently-fitted background-PNP 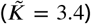, and whilst the parameter estimate for *K* under globally-fitted inertial-PNP (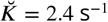; 95% bootstrapped CI: 0.6, 3.6 s^−1^) is a little higher than the median value of the parameter estimate for *K* under independently-fitted inertial-PNP 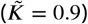, the confidence interval of the former nevertheless contains the latter. Finally, when fitted globally to all flights, both PNP models have shorter optimal delays (inertial-PNP: 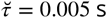 95% bootstrapped CI: 0, 0.03 s; background-PNP: 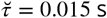; 95% bootstrapped CI: 0, 0.045 s) than when those delays are fitted independently to each flight (see Tables 2 and 3).

Related to these results, it is clear by inspection of Figure 5 that the goodness of fit of the globally-fitted guidance models is more sensitive to the value of *K* for background-PNP than for inertial-PNP. That is, whereas 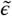 is relatively insensitive to *K* over much of the range plotted for inertial-PNP (i.e. the slope 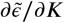 is small in Figure 5A,C), the median RMS error 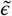 decreases rapidly with increasing *K* for background-PNP at low values of *K* (i.e. the slope 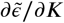 is strongly negative in Figure 5D,F). This presumably explains why the bootstrapped confidence interval for 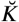 is tighter for background-PNP (95% bootstrapped CI: 3.0, 3.8 s^−1^) than for inertial-PNP (95% bootstrapped CI: 0.6, 3.6 s^−1^).

In summary, although both globally-fitted versions of the PNP guidance law fit the steering of Harris’ hawk attack trajectories comparably well, the best-fitting values of their three guidance parameters are identified with greater certainty for background-PNP than for inertial-PNP. However, globally-fitted inertial-PN fits the data almost as closely as inertial-PNP and globally-fitted background PNP, so provides a more parsimonious model of the data with only two fitted guidance parameters. As an intuitive graphical summary of the model respective fits to the data, we compare the measured trajectories to those simulated under each globally-fitted model, for sample flights corresponding to the first, second, and third quartiles of the RMS error for each model (Figure 6). Given that PP and background-PN perform much more poorly by comparison, we carry forward only the globally-fitted inertial-PNP, background-PNP, and inertial-PN guidance models for formal model validation.

**Figure 6.**
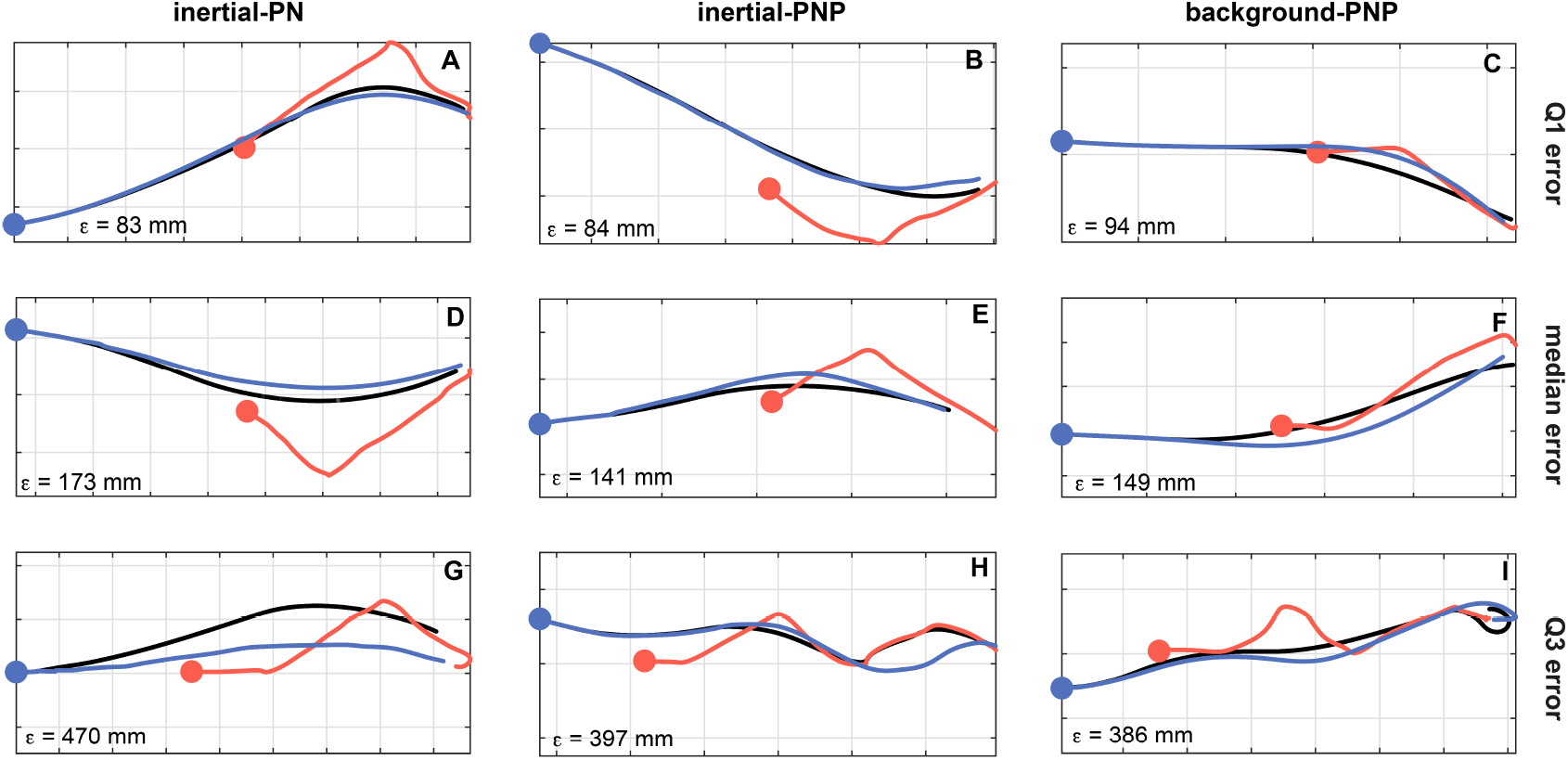
Simulated trajectories for nine sample flights under globally-fitted inertial-PN, inertial-PNP, and background-PNP. Given the measured trajectories of the hawk (blue) and lure (red), we generate the hawk’s simulated trajectory (black) by solving for the model parameters that minimise the median RMS error 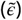 between the measured and simulated trajectories across all n=228 flights. Simulations are shown for: **(A,D,G)** inertial-PN at 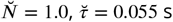; **(B,E,H)** inertial-PNP at 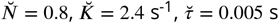; **(C,F,I)** background-PNP at 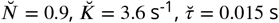; see text for details. Flights are sampled by plotting the simulations for which 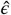 corresponds to the first (A-C), second (D-F) and third (G-I) quartiles of the RMS error *ϵ* for each guidance law. The specific value of 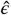 associated with each simulation is shown in the bottom left corner of each panel. Grid size: 1 m

### Model validation

As a final validation step, we apply *k*-fold cross-validation to each of the three globally-fitted guidance models (see Methods and Materials for details). This entails splitting the n=228 flights into *k* = 12 subsets, and applying the global model-fitting approach to the data *k* = 12 times, leaving out a different one of the *k* = 12 subsets as a validation subset. The predictive power of the model is then assessed by computing the median RMS error 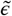 for the validation subset, using the model parameters fitted to the *k* − 1 = 11 subsets. We summarise the results of this validation step by reporting the mean of the median RMS error 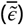 over all *k* = 12 validation subsets for each guidance model. The results of this *k*-fold cross validation show that the order of model predictive ability mirrors the order of model success under the global fitting approach. Specifically, inertial-PNP has the best predictive power 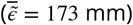, followed by background-PNP 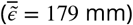, then inertial-PN guidance 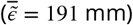. In each case, the mean of the median RMS error for the validation subsets not included in the model fitting 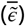 is only 10 − 20% higher than the median RMS error when all of the data are included in the model fitting 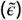, confirming that the approach of fitting the model parameters globally to all flights is successful in avoiding over-fitting to the sample.

## Discussion

The results of our study confirm and extend the results of the previous work on Harris’ hawk pursuit by ***Brighton and Taylor*** (***2019***). Whereas the previous study (***Brighton and Taylor, 2019***) analysed a sample of n=50 flights collected outdoors from N=5 birds using multi-camera videogrammetry, the present study analysed a sample of n=228 flights collected indoors from *N* = 4 birds using motion capture, with one individual shared across both samples. The experimental setup was similar in each case, with an artificial lure being dragged around a set of pulleys producing several unpredictable turns down a zigzag course. We begin by considering the extent to which the present work confirms the conclusions of the earlier study by ***Brighton and Taylor*** (***2019***), before exploring the several ways in which it extends these.

### Harris’ hawk pursuit trajectories are well-modelled by both inertial-PNP and inertial-PN

Consistent with the earlier study by ***Brighton and Taylor*** (***2019***), we found that inertial-PNP modelled our data significantly more closely than inertial-PN when fitted to each flight independently. However, whereas ***Brighton and Taylor*** (***2019***) also found that their globally-fitted inertial-PNP model (with only three parameters in total) fitted their sample of n=50 flights more closely than their independently-fitted inertial-PN model (with one hundred parameters in total), the same result does not hold true for the much larger sample of n=228 flights that we collected here (Tables 2 and 3). Given the obvious risk of over-fitting that exists when fitting the model parameters to each flight independently, we conclude that our data provide no strong statistical reason to favour inertial-PNP over inertial-PN. On the contrary, inertial-PN offers the more parsimonious model of our data if it is assumed that hawks use visual-inertial information to guide their pursuit behaviour (Table 1). The alternative PP guidance law that we and ***Brighton and Taylor*** (***2019***) tested requires only visual information (Table 1), but whereas ***Brighton and Taylor*** (***2019***) found that PP modelled their sample of n=50 flights about as closely as inertial-PN, PP produces a significantly poorer fit than inertial-PN to the n=228 flights that we collected here, regardless of whether the models are fitted to each flight independently (sign test: *p* < 0.0001) or to all flights globally (sign test: *p* = 0.007). We therefore reject PP as an adequate model of Harris’ hawk pursuit behaviour.

The numerical values of the guidance parameters that we have fitted here also corroborate the results of the earlier study on Harris’ hawks by ***Brighton and Taylor*** (***2019***). The parameter estimates display a particularly high degree of consistency for inertial-PN: the median value of *N* fitted to each flight independently was 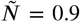 in both studies, whilst the globally-optimised value of *N* was 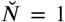 in each case. Such quantitative repeatability is rare in studies of animal behaviour (***Frommlet, 2020***), which makes this a particularly striking result given that significantly higher values of *N* were identified in other studies of peregrine falcons (***Brighton et al., 2017***, ***2021***). Moreover, the parameter setting of *N* = 1 that we have twice identified empirically in Harris’ hawks under inertial-PN is also the value required theoretically to produce the tail-chasing behaviour that is typical of Harris’ hawk pursuit flights (***Brighton and Taylor, 2019***). Conversely, the higher values of *N* that have been identified previously in peregrine falcons (***Brighton et al., 2017***) are necessary to head-off a target during interception. Hence, the parameter estimates that we observe empirically under inertial-PN are not only consistent within a species but also make sense functionally between species.

By contrast, the parameter estimates for *N* and *K* under inertial-PNP are similar, but non-identical, across the present study and the previous work on Harris’ hawks by ***Brighton and Taylor*** (***2019***). Specifically, whereas parameter settings of 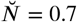 and 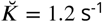 were found to be globally optimal across the n=50 flights modelled by ***Brighton and Taylor*** (***2019***), we found parameter settings of 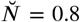 and 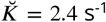 to be optimal across the n=228 flights that we analysed here. Noting that the best-fitting value of *K* is conditional on the setting of the time delay *τ* (Figure 5), it is likely that this difference in the fitted values of *K* between the two studies is coupled to the difference in the fitted time delay, which was shorter in the current study 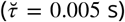 than the previous one 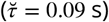. Moreover, it is plausible that this difference in the fitted delay relates to the different measurement technologies used, since the small retroreflective markers used in the previous study were more-or-less rigidly fixed to the body, whereas the larger polystyrene balls that were loosely attached to the birds studied by ***Brighton and Taylor*** (***2019***) could have lagged the body’s motion during turning.

### Background-PNP models hawk pursuit trajectories almost as closely as inertial-PNP

The inertial-PN and inertial-PNP models that we compared in the previous section both require the target line-of-sight 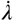 to be measured in an inertial frame of reference, which we denote by writing 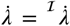. As discussed in the Introduction, the hypothesis that Harris’ hawks implement inertial-PN or inertial-PNP therefore implies involvement of the vestibular system in the guidance loop. In contrast, PP guidance only requires measurement of the target deviation angle *δ* with respect to the pursuer’s velocity vector, which can be done using visual information alone (***Ros and Biewener, 2017***). For instance, the target deviation angle could be measured by comparing the retinal position of the target to the retinal position of the singularity in the optical flow field, which perfectly coincides with the direction of travel during pure translational motion. However, as we have already found that PP does not model our data as closely as inertial-PN or inertial-PNP, visual assessment of the target deviation angle alone is unsupported as an explanation of our hawks’ observed pursuit behaviour.

On the other hand, the background-PN and background-PNP guidance laws that we have defined and tested can also be implemented using visual information alone. This is because they each define the target line-of-sight rate 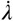 with respect to its visual background, which we denote by writing 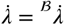. Optimising their guidance parameters globally shows that background-PN provides a very poor model of the data (Tables 3). However, background-PNP performs almost as well as inertial-PNP in modelling our data (Table 3), regardless of whether the model parameters are fitted to each flight independently (sign test: *p* = 0.64; n=228) or to all flights globally (sign test: *p* = 0.55; n=228). Hence, if Harris’ hawks do implement their pursuit guidance using visual information alone, then it follows that they must be implementing background-PNP rather than PP or background-PN amongst the set of five candidate models that we have tested (Table 1).

Noting that background-PNP is just the linear sum of PP and background-PN, these findings lead to three fundamental questions. First, what sensory information is gained by combining measurements of target deviation angle *δ* and background line-of-sight rate 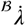 in background-PNP? Second, how does this sensory information compare to the sensory information that is obtained by combining measurements of target deviation angle *δ* and inertial line-of-sight rate 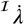 in inertial-PNP? Third, given the superficial similarity between background-PNP and inertial-PNP, how and why do their respective guidance parameters differ as we have found (see Results)? We answer each of these questions in the next section, through an in-depth analysis of the various guidance laws.

### A functional comparison of the different guidance laws

To understand why background-PN explains our data so poorly, whereas background-PNP explains it so well, it is helpful to begin by considering why inertial-PN produces an effective tail-chase at the best-fitting parameter value of 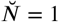. It is clear by inspection of the inertial-PN guidance law (Table 1), that setting *N* = 1 simply equates the pursuer’s inertial turn rate to the target’s delayed inertial line-of-sight rate. Hence, if the pursuer’s initial velocity is directed at the target, then inertial-PN at *N* = 1 will keep the pursuer’s velocity directed at the target if the delay is small. This will serve to correct any target motion across the line-of-sight, thereby commanding a tail-chase. The same holds true if we replace the inertial line-of-sight rate 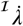 with the background line-of-sight rate 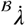, because the change in direction of the line-of-sight resulting from the target’s own motion is independent of whether it is measured with respect to an inertial frame of reference or the visual background. This being so, why does background-PN perform so much worse than inertial-PN in modelling our data?

The reason for this can be understood by comparing the behaviour of inertial-PN and background-PN when the pursuer’s velocity is not directed at the target initially. For clarity, we will also assume that the target remains stationary, although this is not necessary to the argument. Under these circumstances, the component of the pursuer’s motion across the line-of-sight will naturally cause a change in the direction of the line-of-sight to the target. Furthermore, it will cause the same change in the direction of the line-of-sight to any background feature falling close behind the target (i.e. any background feature whose distance to the target is negligibly small). It follows that whereas the change in direction of the target line-of-sight will be registered fully by inertial-PN feeding back the inertial line-of-sight rate 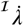, it will not be registered at all by background-PN feeding back the background line-of-sight rate 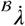. This is because the apparent motion of the target due to the pursuer’s own self-motion will be cancelled by the apparent motion of the background against which it is measured. Hence, whereas inertial-PN will tend to correct for any initial misdirection of the pursuer’s velocity vector in the same way that it corrects for actual target motion, background-PN has no such tendency (see Supplementary Text).

It follows that background-PN will be ineffective at dealing with any difference in the directions of the pursuer’s velocity vector and the target’s line-of-sight that may exist at the outset of the pursuit or that may subsequently be introduced by any error or delay in the pursuer’s response to target motion. Since the difference in these directions is just the deviation angle *δ*, this result also explains why feeding back the deviation angle *δ* in addition to the background line-of-sight rate 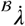 makes background-PNP successful in producing a tail-chase from any set of initial conditions. The same reasoning can also be used to explain the observed similarities and differences in the numerical values of the guidance parameters fitted under background-PNP and inertial-PNP. Specifically, to produce the tail-chasing behaviour that is observed in Harris’ hawks, the component of the line-of-sight rate which results from the target’s motion must be fed back with unity gain, which explains why the best-fitting value of the guidance parameter *N* is 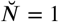 under both background-PNP and inertial-PNP. However, in the case of inertial-PNP, this line-of-sight rate feedback will also have the effect of correcting for any changes in the line-of-sight direction that result from the pursuer’s own self-motion. We should therefore expect that the deviation angle will need to be fed back with lower guidance gain under inertial-PNP than under background-PNP (see Supplementary Text), which is what we observe when comparing the best-fitting values of the guidance constant *K*. Specifically, the guidance gain associated with this deviation angle feedback takes the value 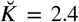 under inertial-PNP and 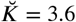 under background-PNP.

### Mechanisation of the three respective guidance laws

The five guidance models that we have fitted to our data (Table 1) are phenomenological models of the bird’s guidance and control. Nonetheless, the practical implementation of each implies a different underlying physiological mechanism. Specifically, the inputs to these guidance laws each entail a different mechanisation, which we illustrate by replacing the black box of each input-output relationship (Table 1) with a block diagram describing the hypothesised information flow (Figure 7). It is important to note that these block diagrams are not unique, in the sense that there may be many different ways of implementing the same input-output relationship. Hence, in the following sections, we discuss how inertial-PN, inertial-PNP, and background-PNP may each be mechanised with reference to the hypothetical block diagrams in Figure 7.

**Figure 7.**
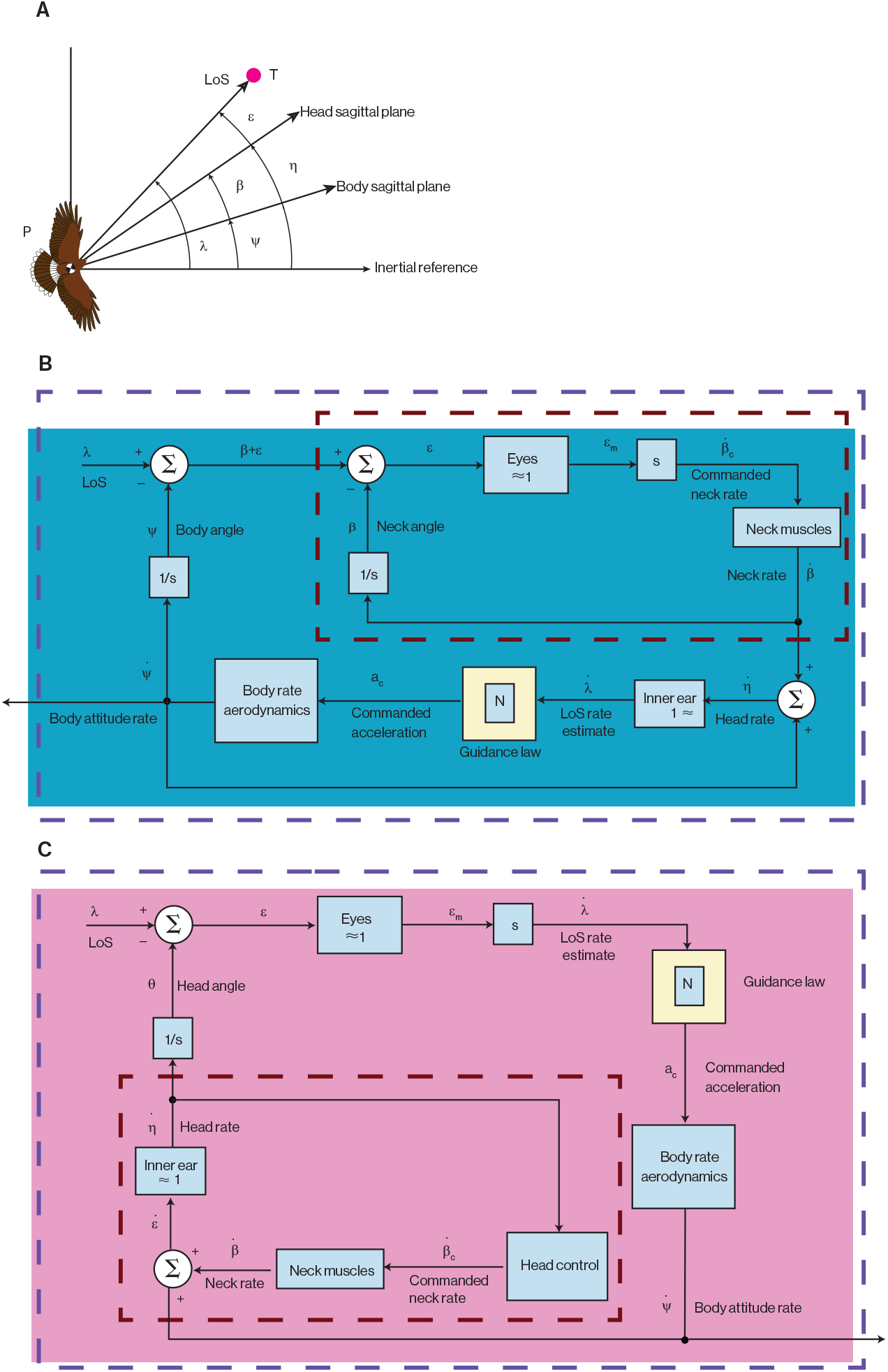
Block diagrams illustrating alternative ways of mechanising inertial-PN. **A** Geometry relevant to the block diagrams. The line-of-sight angle, *λ*, defined in an inertial frame of reference, is a composite of the body angle (*ψ*), the neck angle (*β*), and an error angle (*ε*). The body angle, *ψ*, is defined as the angle between the inertial reference direction and the hawk’s longitudinal body axis, whereas the neck angle, *β*, is defined as the angle between the longitudinal body axis and the sagittal plane of the head. The error angle, *ε*, is defined as the angle between the head sagittal plane and the target azimuth in retinal coordinates. Note that the head angle, η is defined as *ψ* + *β*. The inertial line-of-sight rate, 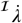, can be reconstructed in different ways. The block diagram in **B** mechanises the estimation of 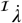 by measuring the angular rate of the hawk’s head, 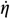, as might the semi-circular canals of the bird’s vestibular system. Here, the visual system drives the inner tracking loop (red dashed line) commanding the neck rate, 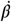, so as to minimise the error between the target and some fixed position on the retina. The vestibular system’s estimate of 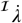 drives the outer guidance loop (blue dashed line) that commands lateral accelerations to control the body attitude rate, 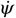, and hawk steering. In contrast, the alternative mechanisation of inertial-PN shown in **C** estimates 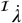 visually from the retinal drift of the target, which it uses to drive the outer guidance loop. For this to provide an accurate estimate of 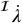, the head must be rotationally stabilised such that 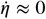. In this case, the vestibular system drives the inner stabilization loop (red dashed line) by commanding 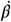 to drive 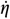 to zero. This figure is adapted from ***Palumbo et al***. (***2010***); other ways of mechanising inertial-PN are possible (see text for details).

#### Mechanisation of inertial-PN

In order to steer an attack trajectory by implementing inertial-PN, a hawk would first need to estimate the inertial line-of-sight rate 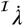 of its target. In homing missiles, this task is usually achieved by a gimbaled seeker that tracks its target by aiming to minimise the target’s drift across its sensor (***Shneydor, 2011***). A sensor called a rate gyro that measures the angular velocity of the seeker can therefore be used to estimate 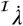 directly from the seeker’s motion (***Murtaugh and Criel, 1966***). Although the details of the implementation may be somewhat more involved in the presence of sensor error and tracking delay (***Palumbo et al., 2010***), it follows that in guided missiles *“*there is no PN without a gyro*”* (***Shneydor, 2011***). If a hawk were to track its target by turning its head like a missile seeker, then the obvious candidates for sensing 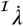 would be the hair cells associated with the semi-circular canals of its vestibular system.

Some authors have argued that these are only able to transduce angular acceleration (***Warrick, 2002***; ***Pennycuick, 2008***), which would have to be integrated to estimate the line-of-sight rate, thereby introducing drift and an unknown constant of integration. However, detailed modelling has suggested that the semi-circular canals may also respond directly to angular velocity (***Mayne, 1950***; ***Goldberg and Fernández, 1975***; ***Muller, 2020***), which would make them better suited to estimating the inertial line-of-sight rate. Either way, under this hypothetical mechanisation of inertial-PN, the visual system would drive the inner tracking loop stabilising the target’s position in the head’s coordinate system (***Hardcastle and Krapp, 2016***), whereas the vestibular system would provide the information on the inertial line-of-sight rate needed to drive the outer guidance loop (Figure 7B). This hypothesis assumes that the eyes move minimally with respect to the head in flight, as is likely to be true for raptors (***Steinbach and Money, 1973***; ***O’Rourke et al., 2010***).

Tracking the target continuously with an imaging device equipped with a gyroscopic sensor is not the only way to measure a target’s inertial line-of-sight rate. For instance, in so-called strapdown missile guidance, the seeker is fixed to the missile’s body (***Palumbo et al., 2010***) such that the drift of the target within the seeker’s coordinate system is just the difference between the inertial line-of-sight rate and the inertial turn rate of the body. In this case, the inertial-line-of-sight rate can be estimated by comparing the rate at which the target drifts across the sensor’s coordinate system to the angular rate of the body. Whilst it is possible that birds use their lumbosacral organ to measure the angular rate of their body (***Stanchak et al., 2020***), it seems unlikely that they use strapdown guidance, because birds do not hold usually their heads fixed with respect to their body (but see ***Tucker*** (***2015***) for a hypothesis to the contrary).

Under yet another alternative mechanisation of inertial-PN, a bird could isolate the movement of its head from that of its body, and keep its head rotationally stabilised even as the body turns. Under this hypothetical mechanisation of inertial-PN, the vestibular system would only drive the inner stabilization loop, whereas the retinal drift of the target would provide the information on line-of-sight rate needed to drive the outer guidance loop (Figure 7C). This hypothesis again assumes that the eyes move minimally with respect to the head, but requires periods of stabilized gaze to be punctuated by fast saccadic head movements, to avoid the target drifting too far across the retina when the head is held stable. Distinguishing between these hypothetical alternative mechanisations of inertial-PN requires detailed quantitative information on head movements during prey pursuit, which will be provided elsewhere.

In the meantime, data collected using head-mounted video cameras suggests the use of both kinds of gaze strategy in birds. Peregrines and other falcons *Falco spp*. have been found to intersperse longer periods of stabilized gaze with short saccadic head movements during the mid-phase of a pursuit, but fixate their target at a consistent location on their retina during its terminal phase (***Kane and Zamani, 2014***; ***Kane et al., 2015***; ***Walker, 2018***). Target fixation has also been observed in a northern goshawk *Accipiter gentilis* chasing terrestrial prey (***Kane et al., 2015***), but unpublished data from Harris’ hawks ***Walker*** (***2018***) suggests that they lie at neither extreme of the mechanisation continuum. Instead, Harris’ hawks allow their target to drift a short distance across the retina, before making a fast saccadic motion to re-fix the retinal coordinates of the target. Comparable variation exists in the target tracking behaviours of different groups of insect. For instance, robber flies (Diptera: Asilidae) stabilise their head rotationally between saccades, whereas dragonflies (Odonata) use their head to track their target (***Mischiati et al., 2015***; ***Lin and Leonardo, 2017***).

#### Mechanisation of inertial-PNP

Implementing inertial-PNP additionally requires measurement of the deviation angle *δ* between the line-of-sight to the target and the pursuer’s velocity vector. The latter’s direction can be estimated directly from the optical flow experienced during translational motion, provided the background is close enough for the pursuer’s self-motion to produce appreciable image motion on the retina. This may explain why evidence for the use of inertial-PNP has so far only been forthcoming in hawks chasing terrestrial targets (***Brighton and Taylor, 2019***) and not in falcons chasing aerial targets (***Brighton et al., 2021***), which instead appear to implement inertial-PN (***Brighton et al., 2017***). The direction of the pursuer’s velocity vector will be most reliably measured if the head is stabilized rotationally, because then the singularity representing the centre of expansion of the optic flow field will coincide with the direction of the velocity vector, which is not true in general when translational and rotational self-motion are combined.

Of course, there are other ways in which a pursuer could estimate the direction of its velocity vector. For example feather airflow sensors could be used to align the bird’s head to the direction of the oncoming air velocity. This will coincide with the bird’s flight direction under still conditions, such that the retinal position of its target would then provide an estimate of the deviation angle *δ*. An even simpler method of estimating the deviation angle would be to approximate the direction of the pursuer’s velocity vector with the direction of its long body axis, since the two are likely to coincide quite closely under still conditions. In this case, if the head were to track the target closely, then the deviation angle *δ* would simply be approximated by the neck angle of the head with respect to the body.

#### Mechanisation of background-PNP

In fact, as we have shown above, measuring the inertial line-of-sight rate 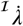 directly need not be necessary if the pursuer were capable of sensing the background line-of-sight rate 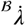 of its target visually. For example, an eagle soaring at great height, or a falcon engaged in an aerial chase against far-off hills, could use the distant background as an inertial reference against which to measure any change in the direction of the line-of-sight to its target. In this case the background line-of-sight rate will approximate the inertial line-of-sight rate. On the other hand, for a Harris’ hawk flying low over the ground whilst viewing its terrestrial prey, the nearby visual background cannot serve as an inertial reference. In this case, the background line-of-sight rate will not approximate the inertial line-of-sight rate, but can nevertheless be used as feedback to the background-PNP guidance law that we have defined above.

Background-PNP requires different mechanisation to inertial-PNP, because the background line-of-sight rate 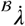 is measured using vision alone. Moreover, measuring 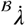 does not even require the head to track the target closely, so the gaze strategy that the bird can use is flexible. This is because the optical flow induced by head rotation is the same for the target and for any background features behind it. It follows that the target’s motion relative to the background is the same independent of any head rotation, which makes background-PNP more robust to flapping perturbations and variation in active gaze than inertial-PNP. On the other hand, background-PNP still requires accurate measurement of the deviation angle *δ*, which is easiest to measure during periods in which the head is rotationally stabilized (see above).

There is one significant limitation of background-PNP, however, which is that for the same physical motion of the target and its pursuer, the target’s apparent motion relative to the visual background will vary depending on the distance between the background and the target. This situation arises because of the distance-dependence of translational optical flow, and creates an ambiguity in the meaning of the background line-of-sight rate. In the context of aerial pursuit of a terrestrial target, however, the distance between the target and its local background is likely to remain approximately constant. Terrestrial pursuit is therefore a best-case scenario for guiding pursuit using background-PNP, as too is pursuit against a background that is su*Z*ciently distant that it can effectively serve as an inertial reference (see above). Either way, the important point is that background-PNP can be implemented using visual cues alone, without requiring the involvement of inertial sensors in the guidance loop.

This conclusion may be especially significant to understanding how insects intercept prey, because the best-known gyroscopic sensors of insects (i.e. the halteres of flies) are located on the body rather than the head. Furthermore, whilst the antennae of hawkmoths (Lepidoptera: Sphingidae) have been found to function as rate gyros (***Daniel et al., 2012***), this function has not yet been confirmed in any predatory insect. In any case, given how well background-PNP models the data, it follows that the use of inertial cues is not strictly necessary to explain the steering behaviour of Harris’ hawks pursuing terrestrial targets. This is not to say that inertial cues will not be important to gaze stabilization, as inertial cues form the basis of the vestibulo-ocular reflex that stabilizes the eyes of birds against their head movements (***Gioanni, 1988***). Rather, we conclude that inertial cues are not necessary to mechanising the guidance loop in background-PNP. Even so, this still begs the question of how visual measurement of the background line-of-sight rate might be mechanised physiologically.

The tectofugal system of a bird brain’s contains certain directionally-selective tectal neurons that respond only to small object motion (***Jassik-Gerschenfeld and Guichard, 1972***), but whose response is strongly modulated by any background motion (***Frost and DiFranco, 1976***; ***Frost et al., 1990***). In particular, their response to small target motion is inhibited by syndirectional background motion and enhanced by contradirectional background motion (***Frost, 1978***; ***Frost et al., 1981***). These tectal neurons therefore respond not merely to retinal drift of a target, but rather to retinal drift of the target relative to its background. In principle, their output may therefore encode the background line-of-sight rate 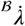 that we have defined. Tectal neuron responses are best known from pigeons, which do not engage in prey pursuit, but neurons with similar responses have also been found in many predatory insects. These include small target motion detectors (STMDs) (***Nordström et al. 2006***; ***Evans et al., 2022***) which synapse with target sensitive descending neurons (TSDNs) whose output commands thoracic muscle activity (***Namiki et al., 2018***). The STMD responses of hover flies (Diptera: Syrphidae) and dragonflies are unaffected by background motion (***Nordström et al., 2006***; ***Evans et al., 2022***), but the TSDN responses of hover flies and robber flies are strongly affected, appearing to depend specifically upon the motion of the target relative to its background (***Nicholas et al., 2018***; ***Nicholas and Nordström, 2021***). Indeed, TSDN response is suppressed completely when the target and the background are moving at the same apparent velocity (***Nicholas et al., 2018***), and this inhibition persists even when the region of syndirectional optic flow is confined to the small area of the frontal visual field where tracked targets are positioned on the retina (***Nicholas and Nordström, 2021***).

Optic-flow sensitive neurons that may be capable of providing background line-of-sight rate feedback are therefore present in both birds and insects, although it remains to be confirmed whether the information that they provide is used as feedback to their guidance loop or simply as a means of enhancing the reliability of target detection.

## Conclusion

To conclude, we find that inertial-PN performs comparably well to the more sophisticated inertial-PNP and background-PNP guidance laws that we have used to model the n=228 Harris’ hawk attack 587 trajectories reported here. Inertial-PN therefore provides an effective framework for modelling the guidance of aerial pursuit behaviour across a broad phylogenetic range of predatory taxa. Unlike the background-PNP model, which requires only visual information to implement guidance, the inertial-PN and inertial-PNP models requires fusion of visual and inertial information. Visual-only pursuit is also possible under the PP guidance law, but PP does not model the data as successfully as inertial-PN. We are left with the conclusion that inertial-PN provides the most parsimonious model of the data that entails fusion of visual and inertial information, whereas background-PNP is the only well-fitting candidate for guidance using visual-only cues. In contrast, inertial-PNP provides the closest fit to the data overall, albeit that this difference is only statistically significant when fitting the model parameters to each flight independently, which almost certainly results in over-fitting. Testing between the different ways in which these guidance laws may be mechanised requires detailed quantitative information on head movements during pursuit, which we will provide elsewhere. However, of particular relevance to the background-PNP model, certain target-detecting neurons of insects and birds are already known to respond to the movement of a target relative to its background, and are therefore sensitive to the background line-of-sight rate that would need to be fed back to command turning under background-PNP.

## Methods and Materials

### Experimental protocol

We flew N=4 captive-bred Harris’ hawks *Parabuteo unicinctus* in pursuit of an unpredictably moving artificial lure indoors. The sample comprised one 7-year old female, which had also been included in the previous study of pursuit behaviour by ***Brighton and Taylor*** (***2019***), plus three first-year males that had not previously chased a target. We conducted the flight trials at the John Krebs Field Station, Wytham, Oxford, UK, in a flight hall with floor dimensions of 20.2 m × 6.1 m, and a minimum ceiling height of 3.8 m. We lit the windowless flight hall using flicker-free LED uplights providing approximately 1000 lux of diffuse overhead light, mimicking overcast morning or evening light levels. The floor of the flight hall was carpeted with artificial grass, and the walls of the flight hall were hung with camouflage netting to provide visual contrast. This work was approved by the Animal Welfare and Ethical Review Board of the Department of Zoology, University of Oxford, in accordance with University policy on the use of protected animals for scientific research, permit no. APA/1/5/ZOO/NASPA, and was considered not to pose any significant risk of causing pain, suffering, damage or lasting harm to the animals.

The hawks performed 157 training flights between January and February 2018 and, from February to March 2018, we flew the same individuals for a total of 330 test flights of which n=228 were retained in the final analysis (see below for details of exclusions). On each trial, the lure was pulled along an unpredictable, zigzagging course across the floor by a tow line attached to two parallel Aerotech ACT140DL linear actuators (Aerotech Limited, Hampshire, UK), as shown in Figure 8A-B. We randomised the lure’s starting position, trajectory, and speed (6 to 8 m s^−1^) across trials, to prevent the hawks learning the target’s motion. The tow line was guided around a system of pulleys, with dummy lines laid along the alternative routes that were not connected to the motor, to avoid giving the hawks any information on what course the lure would follow until it began moving. The lure was hidden inside a tunnel, out of sight of the hawk, at the start of each trial. This arrangement was designed to simulate the evasive behaviour of the lagomorph prey that Harris’ hawks hunt, which jink sharply from side to side when evading predators (***Bednarz, 1988***).

**Figure 8.**
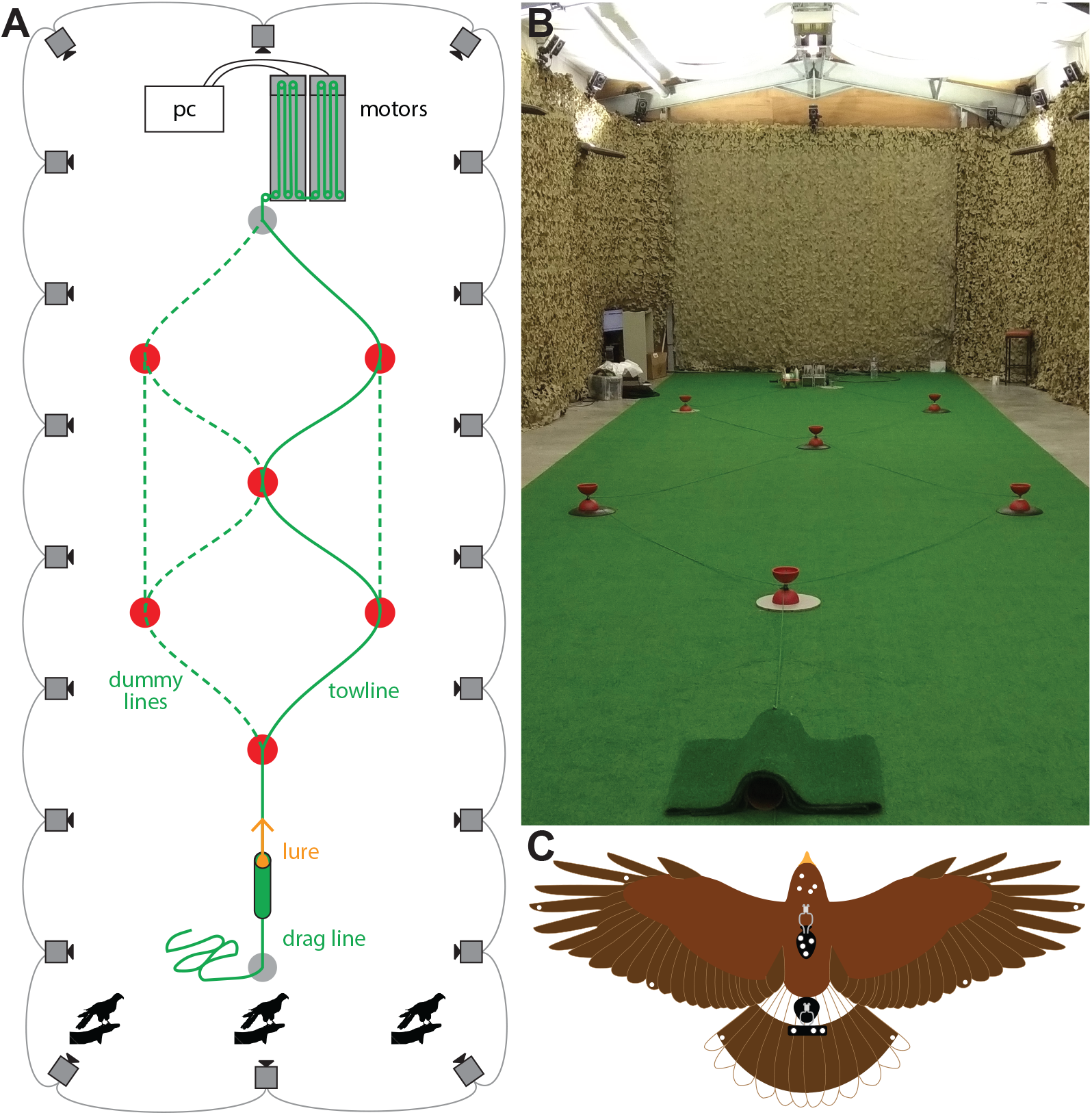
The setup of the pursuit task. **(A)** Schematic of experimental setup from an overhead view. The approximate positions of the 20 motion capture cameras are indicated by the camera icons, while the bird icons indicate the three alternative starting positions for the bird. The approximate positions of the 6 pulleys are shown as red circles, and the 6 alternative lure paths are shown as green lines. To prevent the bird from predicting the lure’s path, we laid dummy tow lines for each trial (dotted green line), in addition to the motor-connected tow line (solid green line). We randomised the lure path between trials. **(B)** Photograph of the set-up, from the perspective of the middle starting position. **(C)** Schematic drawing of a Harris’ hawk wearing the rigid templates (black) to which we attached the spherical retroreflective motion capture markers (white circles).

### Motion capture

We recorded the flight trajectories using twenty motion capture cameras (Vicon Vantage 16, Vicon Motion Systems Ltd, Oxford, UK) positioned 3 m above floor level around the walls of the flight hall. The motion capture cameras sampled the positions of the markers at 200 Hz under stroboscopic 850 nm infrared illumination. We calibrated the cameras using an Active Calibration Wand (Vicon Motion Systems Ltd, Oxford, UK) that we also used to set the capture volume origin and ground plane. To record the flight trajectories, we fitted each hawk with two rigid templates, each holding several 6.4 mm spherical retroreflective markers (Figure 8C). We attached one template holding four asymmetrically arranged markers to the back using a falconry harness (TrackPack Mounting System, Marshall Radio Telemetry, UT, USA), and another template holding three symmetrically arranged markers to the tail using a falconry tail mount (Marshall Aluminium Tail Feather Piece, Marshall Radio Telemetry Ltd, Cumbria, UK). We also attached 3 mm markers to the head, wings, and tail, although we do not analyse these data here. To record the lure trajectory, we attached three 6.4 mm markers to each long face of the lure. We also attached markers to the pulley system, and the tunnel from which the lure started.

### Data processing

We reconstructed each marker’s position with respect to a coordinate system fixed to the principal axes of the flight hall, using Vicon Nexus 2.7.6 software (Vicon Motion Systems Ltd, Oxford,UK). However, as the Vicon software proved unreliable in labelling the markers between frames (i.e. identifying the same marker from one frame to the next), we wrote a custom marker labelling script in MATLAB R2019a (The Mathworks Inc., MA, USA). This script first pooled those markers that remained within the same 10 mm range for ≥ 60% of the frames and labelled them as stationary markers; these are markers on the pulley system and starting tunnels. By a process of exclusion, the remaining unlabelled markers are therefore either bird or lure markers. We identified lure markers as any markers falling within 150 mm of the floor plane, and identified the remaining markers as bird markers. These bird markers include single markers placed directly on the head, wings, and tail, in addition to the several markers fixed rigidly to the two templates. We therefore used the pairwise distances between markers on the two rigid templates to define, search for, and label them. Due to wear or damage, we had to replace the markers occasionally, and therefore defined the pairwise distances for each template on a per-day basis. The pairwise distances between each of the markers in the same frame were compared against, and matched to, each template with a tolerance of 5 mm in order to label the backpack and tail mount markers correctly (see also (***KleinHeerenbrink et al., 2022***)).

To avoid including flights with excessive data dropout resulting from marker occlusion, we only carried forward those flights in which the back template markers could be reconstructed in > 50% of the recorded frames. This left us with a reduced sample of 274 flights as candidates for further analysis. We used the centroid of the backpack markers, tail-mount markers, and lure markers as an initial estimate of the position of each object. To enable reliable reconstruction of each centroid, we considered any frames in which fewer than 3 markers were detected for either the template or the lure as missing data. The remaining trajectory data nevertheless contained some misidentified markers, which we removed from the raw data using a two-step process of mean sliding window elimination. In a first step, we removed any extreme outliers, defined as points falling > 500 mm from the centroid trajectories smoothed using a 0.05 s window. Having removed these extreme outliers, we then applied the same method in a second more conservative pass to eliminate moderate outliers at a distance threshold of > 75 mm.

After excluding outliers from the raw data streams for the bird and lure in this way, we cropped each sequence to begin at the first frame for which both the bird and lure were visible. We then used the centroid of the identified backpack and tail-mount markers to define the position of the bird, and used the centroid of the identified lure markers to define the position of the lure. Of the 274 flights remaining after excluding any with excessive data dropout (see above), we only analysed those flights in which the hawk successfully intercepted the lure, which we determined according to whether the hawk came within 30 mm of the lure. Having applied this stringent selection criterion, we were left with a total of n=228 flights that we took forward to the main guidance analysis. We used cubic interpolation to fill in missing data points and fitted a quintic spline to smooth the data, using a spline tolerance of 0.03 m for the bird data and 0.01 m for the lure data. Finally, we differentiated and evaluated the splines analytically in order to estimate the velocity and acceleration of the bird and lure at 20 kHz, as required to ensure a reasonably small integration step size for the forward-Euler method simulations that we describe in the next section.

### Simulations of guidance behaviour

Because the measured pursuits all occurred just above the ground plane, we only model the horizontal components of the trajectory data in our simulations. We simulated the hawk’s flight trajectory by using each of the different guidance laws in Table 1 to model its predicted turn rate. We used the hawk’s measured position and velocity to define the initial conditions for the simulation, and forced the hawk’s simulated flight speed to match its measured flight speed. We used the measured time-history of the lure’s position and velocity to compute the target’s deviation angle *δ*, inertial line-of-sight rate 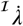, and background line-of-sight rate 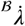, feeding back the relevant quantities to command turning.

The line-of-sight vector ***r*** is defined as:

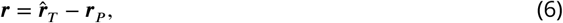

where 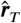 is the measured position of the target and ***r_P_*** is the simulated position vector of the hawk. We calculate the scalar deviation angle *δ* as:

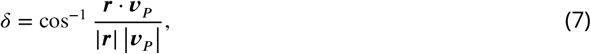

where ***v_p_*** is the simulated ground velocity of the hawk. For the purposes of the guidance modelling, the deviation angle *δ* is more conveniently represented in vector form as:

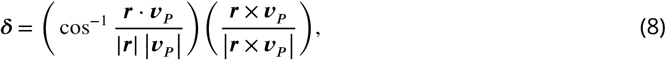

The angular velocity of the line-of-sight vector ***r*** depends on the frame of reference in which its time derivative is taken. Specifically, the inertial line-of-sight rate 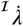 is calculated in vector form as:

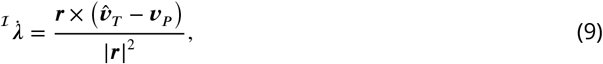

in which 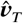 is the measured ground velocity of the target, whereas the background line-of-sight rate 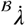 is calculated in vector form as:

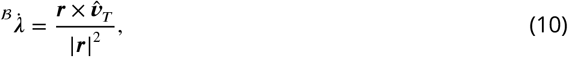

which we derive in the Supplementary Text.

Each of the five guidance models produces turning at a rate 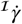 by commanding a centripetal acceleration ***a***. The five guidance laws displayed in Table 1 produce turning at a rate 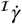 by commanding a centripetal acceleration ***a***. In particular, inertial-PNP commands this acceleration as:

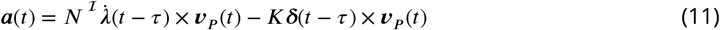

whereas background-PNP commands acceleration as:

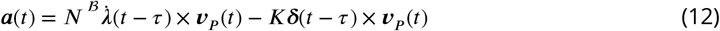

in which *τ* is a fixed sensorimotor time delay. PP guidance is produced by setting *N* = 0 in either case, whereas inertial-PN and background-PN are produced by setting *K* = 0 in the equations for inertial-PNP and background-PNP.

We simulated the hawks’ trajectories under each of these five guidance laws by implementing the preceding continuous time equations in discrete time in MATLAB, coupling them with the following pair of difference equations:

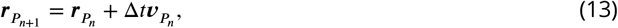

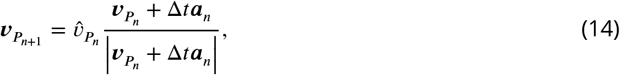

in which 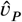 is the hawk’s measured ground speed, and where the subscript notation indicates the values of the variables at successive time steps such that *t*_*n*+1_ = *t_n_* + Δ*t*.

### Model fitting

We optimised the parameter values of each guidance model by simulating each hawk trajectory independently under different values of the time delay *τ*. We varied *τ* on the interval 0 ≤ *τ* ≤ *τ_max_*, where *τ_max_* = 0.15 s, at increments corresponding to the 0.005 s sampling period of the Vicon system. Taking time *t* = 0 as the point at which the lure began moving, we simulated the bird’s flight trajectory from *t* = *τ_max_* to the point of intercept as defined above. This was done to ensure that we always simulated the same section of data for each flight. At each value of *τ*, we used a Nelder-Mead simplex algorithm to find the values of *N* and/or *K* that minimised the root mean-square (RMS) error ε between the simulated (***r_p_***) and measured 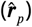 positions of the bird:

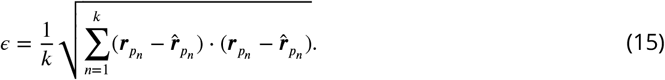

Using this approach, we found the best-fitting values of the parameters *τ*, *N* and/or *K* for each flight and for each guidance law independently, together with the associated value of the RMS error *ϵ*.

In the Results, we report the medians of these parameters over all the flights as 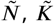, and 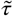, together with the corresponding median RMS error 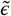. We also fitted the inertial-PNP and background-PNP guidance laws globally across all of the flights, noting that these models both contain PP as a special case when *N* = 0, and each contain inertial-PN and background-PN as special cases when *K* = 0. Specifically, we found the unique combination of *N*, *K* and *τ* that minimised the median RMS error 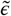 by varying *N* on the interval [0, 1.5] at 0.1 increments, *K* on the interval [0, 5] s^−1^ at 0.2 s^−1^ increments, and *τ* on the interval [0, 0.15] s at 0.005 s increments. In the Results, we report the globally optimal values of these parameters over all the flights as 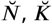, and 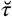, together with the corresponding median RMS error 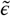. It is important to note that our minimization of the RMS error (i.e. L2-norm) here differs from our minimization of the mean absolute error (i.e. L1-norm) in a complementary analysis of these data in the context of obstructed pursuit. (***Brighton et al., 2023***). The L2 norm penalizes larger deviations to a greater extent than smaller deviations, so the use of the L1 norm was more robust to perturbations to the pursuit trajectories caused by the presence of obstacles in the latter study.

### Guidance model comparison

We computed bootstrapped 95% confidence intervals (CIs) for the median best-fitting parameter estimates 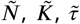 and 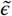, using the bias-corrected and accelerated percentile method over 10^6^ iterations in MATLAB. In other words, beginning from the original sample of n=228 flights, we re-sampled with replacement those flights that contributed to the calculation of the median values of the parameters. We also calculated bootstrapped 95% confidence intervals (CIs) for the globally optimal parameter estimates 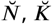, and 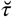, together with the corresponding median RMS error 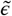. In this case, we used the same bias-corrected and accelerated percentile method, but over 10^3^ iterations. In other words, beginning from the original sample of n=228 flights, we re-sampled with replacement those flights that contributed to the identification of the model parameters at which the median RMS error 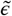 was minimised.

Finally, to assess the predictive power of different guidance models, we performed a *k*-fold cross-validation analysis on the inertial-PN, inertial-PNP, and background-PNP models. Specifically, we first split the n=228 flight trials into *k* = 12 data subsets, each containing 19 flights. We then applied the global fitting technique to *k* − 1 = 11 of the subsets, finding the optimal parameters as those that minimise 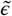. The predictive power of the model parameters is then the 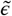 calculated on the remaining single subset. We applied this process 12 times, excluding a different subset each time, and we reported the predictive power of the different models as the mean of the median RMS error 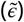 over all *k* = 12 validation subsets given each guidance model.

## People

We thank members of the Oxford Flight Group for insightful comments and helpful discussion. We thank L. Larkman, H. Sanders, and M. Parker for their assistance as falconers during experiments and in husbandry of the Harris’ hawks. We thank H. Frost for comments on the manuscript.

## Author contributions

Conceptualization: J.A.K., C.H.B., and G.K.T.; Formal analysis: J.A.K., C.H.B., L.A.F., M.K.H., J.S., and G.K.T.; Investigation: J.A.K., C.H.B., M.K.H., and S.M.; Methodology: J.A.K., C.H.B., and G.K.T.; Supervision: G.K.T.; Visualization: J.A.K. and C.H.B.; Writing*—*original draft: J.A.K.; Writing*—*review and editing: G.K.T. All authors reviewed and commented on the original draft. Authors other than the first author are listed alphabetically.

## Data and materials availability

Supporting data and code are available at figshare: https://doi.org/10.6084/m9.figshare.21769982.

## Supplementary Text

Here we show that the background line-of-sight rate is approximately equal to the part of the inertial line-of-sight rate that is due to the target’s motion. We express the position of the target (T) and the pursuer (P) using the position vectors ***r_T_***, and ***r_P_***. These position vectors are expressed with respect to the origin *O* of an inertial frame of reference 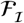 defined by the orthonormal basis vectors {**1**_***x***_, **1**_***y***_, **1**_***ω***_}. The line-of-sight vector from the pursuer to its target is ***r*** = ***r_T_*** − ***r_P_***, and the time derivative of ***r*** with respect to 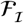 is the relative velocity ***v*** = ***v_T_*** − ***v_P_***. We denote the angular velocity of the line-of-sight vector ***r*** in 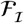 as 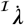, where:

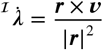

which we may decompose as:

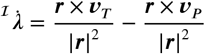

which shows that the target’s line-of-sight rate is composed of a part due to the target’s motion and a part due to the pursuer’s own self-motion.

By analogy, if we express the position vector of some stationary background point (B) in 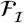 as ***r_B_***, then the angular velocity of the line-of-sight to that background point is just:

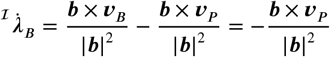

where ***v_B_*** = 0 is the velocity of the background point in the inertial frame of reference, and where ***b*** = ***r_B_*** − ***r_P_*** is the line-of-sight vector to the background point. Hence, if the pursuer measures the apparent motion of its target with respect to this background point, it will measure:

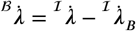

which we may expand as:

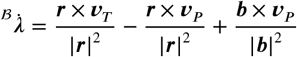

For distant background points, the last term on the right hand side vanishes as ***b*** → ∞, such that 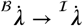 as ***b*** → ∞. However, for the case we consider here, in which the background point is immediately behind the target such that ***b*** ≈ ***r***, we have instead that:

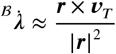

which justifies the formula for 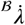 given in Equation 10 of the Methods and Materials, and shows that the background line-of-sight rate is approximately equal to the part of the inertial line-of-sight rate that is due to the target’s motion. Finally, we may take the line-of-sight vector **b** to the stationary background point (B) as defining the direction of a new basis vector 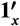 with its origin at (P) as shown in Figure 2. If we define another new basis vector 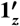 with its origin at (P) that points in the same direction as the basis vector **1**_***z***_ from 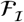, then together with the orthonormal basis vector 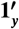, this completes the definition of the background frame of reference 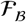 that we refer to in the main text.

These results may be used to formalise the argument made in the Discussion to explain why background-PN is ineffective at steering a pursuit. As we have shown, the background line-of-sight rate 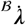 that is fed back under background-PN contains no information on the pursuer’s own self-motion, so cannot be used to correct any initial misdirection of the pursuer’s velocity. On the other hand, the deviation angle *δ* does contain information on the pursuer’s self-motion, since it is defined in vector form as:

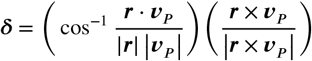

Hence, the reason why background-PNP is effective in steering a pursuit, whereas background-PN is not, is that feeding back information on the deviation angle *δ* reintroduces information on the pursuer’s own self-motion.

The relationship between inertial-PNP and background-PNP can be shown explicitly by using the equations for 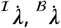, and ***δ*** above to write the inertial-PNP guidance law as:

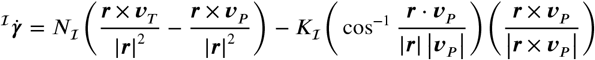

where 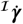 is the pursuer’s turn rate in the inertial frame of reference. Likewise, we may write the background-PNP guidance law as:

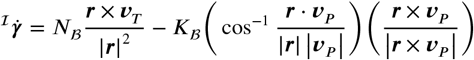

where we have used subscript notation to distinguish the values of the parameters *N* and *K* under the two different guidance laws. Hence, if we assume that *N_I_* ≈ *N_B_* ≈ 1, which is the result we find empirically, then it follows that to command approximately the same observed amount of turning under each guidance law we require:

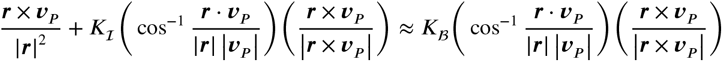

which we may simplify as:

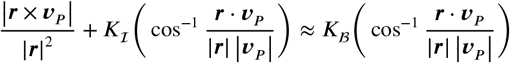

and may then rearrange to yield:

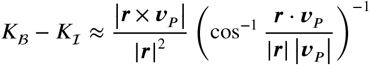

Hence, as the right-hand side of this equation is non-negative, we should expect that *K_B_* > *K_I_*, which is also the result that we find empirically (see Results).

